# Heparan sulfate-function is essential for Integrin-dependent cell-matrix interactions and regulates glycosaminoglycan synthesis synthesis through YAP

**DOI:** 10.64898/2026.01.08.697362

**Authors:** Ann-Christine Severmann, Christoph Waterkamp, Meike Buchholz, Isabel Adorf, Lutz Fleischhauer, Julius Sefkow-Werner, Katja Jochmann, Tatjana Holzer, Velina Bachvarova-Matic, Nina Schulze, Johannes Koch, Bent Brachvogel, Elisa Migliorini, Hauke Clausen-Schaumann, Perihan Nalbant, Daniel Hoffmann, Andrea Vortkamp

## Abstract

Extracellular matrix (ECM) is the main component of cartilage, making it an ideal environment to study cell-matrix interactions. Among ECM constituents, heparan sulfate (HS)-carrying proteoglycans (PGs) are of particular interest since they are not only structural components but are also involved in cell matrix adhesion and signalling processes. We previously demonstrated that transgenic mice with a clonal loss of HS synthesis in chondrocytes (*Col2-rtTA-Cre;Ext1^e2fl/e2f^*^l^) develop clusters of enlarged cells in the articular cartilage (AC), which are surrounded by a glycosaminoglycan (GAG)-rich ECM. This led to the questions how HS regulate the molecular composition and mechanical properties of the ECM, how they sense alterations in the HS structure and how they respond to it.

We stained tissue sections of *Col2-rtTA-Cre;Ext1^e2fl/e2f^* animals and detected increased levels of chondroitin sulfate (CS), Aggrecan (Acan), Perlecan (Pcan), Matrilin (Matn)-3 and-4, Collagen type II (Col2) and Col9, while Col12 was abolished in the HS-deficient clusters. We assessed the stiffness of the mutant matrix by Atomic Force Microscopy (AFM) and found that it was markedly softer than the surrounding, HS-containing tissue. Likely in response to this altered texture, HS-deficient clones showed increased protein levels of Integrin pathway components. To model a loss of HS-function *in vitro*, we treated murine embryonic fibroblasts (MEFs) with the HS-antagonist *Surfen*. Treatment during cell adhesion resulted in impaired cell-substrate adhesion, increased formation of filopodia-like membrane protrusions, decreased cell polarisation and migration, reduced formation of FA and SF, and a translocation of YAP into the cytoplasm. Similarly, we observed reduced cell polarisation in HS-deficient CHO pgsD-667 cells, which could not be rescued by external presentation of HS. When MEFs were treated with *Surfen* after the completion of the initial cell adhesion process, inhibition of HS-function led to an increased formation of FA and SF, in line with the increased levels of Integrin pathway components observed in HS-deficient chondrocytes *in vivo*. We detected high levels of Yes1-associated protein (YAP) in the HS-deficient clusters, and we investigated the effect of YAP modulation on high density micromass cultures from primary murine chondroprogenitors. YAP activation induced an increased GAG synthesis similar to Surfen, while YAP inactivation partially abolished the effect of Surfen, showing that YAP acts downstream of HS function and controls GAG synthesis.

Taken together, we demonstrated that HS-function is essential for Integrin-dependent cell-matrix interactions. Information on the impaired cell matrix adhesion upon loss of HS is conveyed into the nucleus via YAP, which at least partially controls the synthesis of GAGs in chondrocytes.

## Introduction

Cartilage, in contrast to other tissues, is constituted by only 2% cells and primarily contains ECM, posing an excellent model system to study cell-matrix interactions. Cartilage ECM is mainly composed of Collagen fibres, giving it tensile strength, and PGs, providing a high osmotic pressure, drawing water into the tissue and creating an elastic gel (Fox, 2009). Although CSPGs are more abundant (Gahunia, 2020), HSPGs are of particular interest since they are not only structural components but interact with at least 437 proteins identified to date (Rudd, 2017) and modulate the activity of numerous secreted proteins relevant for the progression of OA, including cytokines, proteases and signalling factors (Ravikumar, 2020).

HS synthesis starts with the addition of a tetrasaccharide linker to a serine residue of the PG core protein. Alternating units of N-acetylglucosamine (GlcNAc) and glucuronic acid (GlcA) are then added by a heterodimeric complex formed by the glycosyltransferases Exostosin (Ext) 1 and 2. The growing polysaccharide chain is further modified by N-deacetylation/-sulfation of GlcNAc by members of the N-deacetylase/sulfotransferase family (Ndst1-4), epimerisation of GlcA to IdoA by GlcA-C5-epimerase, 2O-sulfation of IdoA by HS-2O-sulfotransferase (HS2St1) and sulfation in 3O- or 6O-position of GlcNAc by HS-3O- and 6O-sulfotransferases. The expression of HS-modifying enzymes is cell type-dependent and thus a tissue-specific modification pattern is established. Due to the negative charge of their sulfate groups HSPGs interact with positively charged moieties of proteins, such as the Cardin-Weintraub motif found in members of the fibroblast growth factor (FGF) and hedgehog (HH) ((Esko, 2002) (Bulow, 2006) (Poulain, 2015) as well as HS-binding motifs in ECM components, such as Collagens, Laminins and Fibronectin (Delacoux, 1998) (Nielsen, 2001) (Mahalingam, 2007).

We and others (Severmann, 2020) ((Sgariglia, 2013) have previously shown that mice with a clonal deletion of Ext1 in chondrocytes develop clusters of enlarged, HS-deficient chondrocytes in the AC that are surrounded by a dense, PG-rich matrix. *Col2-rtTA-Cre;Ext1^e2fl/e2f^* mice have originally been established to investigate the formation of osteochondromas (Jones, 2010). We demonstrated that the clustered cells found in the AC these animals are not hypertrophic and that the mice are protected from OA (Severmann, 2020) (. Additionally, we reported that reduced HS levels (*Ext1^gt/gt^*) or altered HS sulfation (*HS2St1^-/-^*) lead to a compensatory increase of CS production in embryonic growth plate (GP) cartilage of mouse mutants (Bachvarova, 2020), indicating that chondrocytes sense an altered HS structure and react by upregulating other ECM components.

Integrin dimers serve as major ECM receptors, mechano-sensors and adhesion molecules, and signal through numerous pathways, such as MAPK/ERK and the small GTPases RhoA, Rac1 and Cdc42 (Huveneers, 2009). During cell adhesion, Integrin subunits are clustered into adhesion complexes which mature to FA upon sufficiently stable substrate binding. After initial activation of the Integrin dimers, the adaptor protein Talin connects the receptor to SF, which are contractile fibres formed by Actin and non-muscular Myosin Heavy Chain Type II (NMII). Upon contraction, Talin is stretched, and a Vinculin binding site is revealed. This interaction of Actin, Talin and Vinculin then drives the clustering of additional Integrin dimers and further components into the nascent FA. Focal adhesion kinases (FAKs) oligomerise, activated by auto-phosphorylation, and in turn phosphorylate Src, which then activates Paxillin. Paxillin then stabilises the FA by binding to Vinculin, simultaneously providing a binding site for Actin-polymerising adaptor proteins. Simultaneously, FAK recruits additional Talin proteins, further accelerating FA formation (Legate, 2009) (Wiesner, 2006). FA then serve as a connection between the ECM and the SF of the cytoskeleton, enabling the generation of traction within the cell, which is essential for cell polarisation and migration (Parsons, 2010). Mechanical cues from cytoskeletal tension, cell geometry and ECM stiffness regulate the subcellular localisation and activity of the signalling factor YAP When cells adhere on soft substrates or on small surfaces, YAP is typically found in the cytoplasm, while it translocates to the nucleus when cells adhere to hard surfaces and when high tensile forces are generates within the cell or applied to it. The translation of mechanical cues into the localisation of YAP is transduced via the canonical Hippo as well as non-Hippo pathways. This includes binding of YAP to filamentous actin in the cell periphery via adaptor proteins such as AMOT when few SF are present, phosphorylation of YAP and subsequent degradation, and translocation of YAP into the nucleus upon stretching of nuclear pores. YAP activity then regulates cell proliferation, survival and differentiation (Dupont, 2011) (Heng, 2021).

In this study, we detected a distinct matrix composition around clusters of HS-deficient cells of *Col2-rtTA-Cre;Ext1^e2fl/e2fl^* mice and a clearly decreased stiffness of the mutant matrix. *In vivo*, we found increased protein levels of Integrin pathway components in the HS-deficient clusters. *In vitro*, treatment of MEFs with HS-antagonist Surfen decreased initial cell adhesion, increased the number of cells with filopodia-like protrusions, impaired cell polarisation and migration, reduced formation of FA and SF, and decreased the translocation of YAP to the cytoplasm. In contrast, treatment of adherent cells with Surfen resulted in increased proportions of cells forming FA and SF, similar to our observation *in vivo.* Addition of a YAP activator to micromass cultures resulted in an elevated GAG synthesis, similar to blocking HS function with Surfen. In contrast, YAP inhibition reduced the effect of Surfen. In line with this, *Col2-rtTA-Cre;Ext1^e2fl/e2fl^* animals showed increased protein levels of YAP in the HS-deficient clusters.

Taken together, our results show that HS-function is essential for Integrin-dependent processes, that the effect of HS-deficiency is mediated, at least partially, via YAP and that signalling via YAP controls GAG synthesis.

## Material and Methods

### Mouse husbandry

All animal experiments were carried out according to the institutional guidelines of the University of Duisburg-Essen and approved by the animal welfare officer. Mouse husbandry was approved by the city of Essen (Az: 32-2-11-80-71/348) in accordance with § 11 (1) 1a of the “Tierschutzgesetz”. Work with transgenic animals was approved by the “Bezirksregierung Düsseldorf” (Az: 53.02.01-D-1.55/12) in accordance with § 8 Abs. 4 Satz 2 of the “Gentechnikgesetz”.

In each individually ventilated cage (“Green Line”, TechniPlast) either up to 6 females, 1 breeding pair, or a single male were kept under SPF conditions, with nesting material (Abed) as environmental enrichment. A 12 hour light/dark cycle was applied, and food (Sniff) and water were provided *ad libidum*.

### Transgenic mice

Transgenic *Col2-rtTA-Cre;Ext1^e2fl/e2fl^*[Tg(Col2a1-rtTA,tetO-cre)22Pjro;Ext1^tm1.1Vcs^] (Jones, 2010), R26R-LacZ [Gt(ROSA)26Sor^tm1Sor^] (Soriano, 1999) and *Ext1^gt/+^* [Ext1^Gt(pGT2^™^pfs)064Wcs^] (Mitchell, 2001) mice were maintained on a C57Bl/6J genetic background. C57Bl/6J wildtype mice were purchased from Envigo. *Col2-rtTA-Cre;R26R-LacZ;Ext1^e2fl/e2fl^* compound animals (later termed *Col2-rtTA-Cre;Ext1^e2fl/e2fl^*) were created by crossing the respective mouse strains. Genotyping of transgenic animals was performed on genomic DNA isolated from biopsies using DirectPCR Lysis Reagent (VIAGEN) and the primers listed in Table S1. Isolation of mouse embryos in the last trimester was approved by “Bezirksregierung Düsseldorf” (Az: 81-02.04.2022.A227). For timed pregnancies of *Ext1^gt/+^* and C57Bl/6J mice, the noon of the day of a vaginal plug was considered to be embryonic day 0.5 (E0.5).

### Doxycycline injection and Cre recombination

Breeding of genetically modified mice potentially burdened by osteochondroma formation (Az: 84-02.04.2015.A111, Az: 81-02.04.2023.A127) and the induction of Cre-recombination by doxycycline administration (Az: 84-02.05.20.12.156 and Az: 81-02.04.2017.A507) were approved by the “Landesamt für Natur, Umwelt und Verbraucherschutz (LANUV) Nordrhein-Westfalen”. Cre activity was induced at postnatal day 8 (P8) by peritoneal injection of 80 mg doxycycline per kg body weight (kgbw) into lactating dams as previously described (Severmann, 2020).

### Immunofluorescence staining of tissue sections

In brief, forelimbs of *Col2-rtTA-Cre;R26R-LacZ;Ext1^e2fl/e2fl^* mice were isolated, fixed with 4%PFA for 48-72h at 4°C and decalcified in 25%EDTA at 37°C for 2-3 weeks, embedded into paraffin and sectioned at 7µm thickness.

Staining for Acan, Pcan, Matn-3 and-4, Cartilage Oligomeric Matrix Protein (COMP) and ColII, IX and XII were performed at Bent Brachvogel’s lab at University of Cologne as previously described (Brachvogel, 2013). Immunofluorescence stainings of CS (clone 2B6, detecting the ΔDi-4S neo-epitope), ERK (Cell Signalling Technologies 4695), pFAK (Invitrogen 44624G), FAK (Millipore 05-537), pERK (Santa Cruz Sc136521) and NMII (Cell Signalling Technologies 3403) were performed using suitable AlexaFluor-labelled secondary antibodies (Thermo Fisher). For CS an antibody retrieval with 50mU/L chondroitinase was performed (Hayes, 2008), for ERK with 1000U/ml hyaluronidase (Sigma-Aldrich), and for Acan with 10% formic acid and chondroitinase digestion. To stain Integrin β1 (ItgB1; Millipore mAB1997) and Src (Cell Signalling Technologies 4060), a signal amplification step using a biotinylated secondary antibody (Vector labs BA-1000 or BA-2000) and a fluorescence-labelled streptavidin (Thermo Scientific S32356) was employed, and antigens were retrieved by boiling in 10mM citrate buffer and hyaluronidase digestion. Nuclear staining was performed using 500ng/ml DAPI in ddH_2_O, slides were mounted with Mowiol/DABCO and immunostained sections were imaged using reflected light fluorescence microscopy (AxioObserver 7, Zeiss).

### Atomic Force microscopy (AFM)

Hindlimbs were harvested from 4 weeks (4w) old *Col2-rtTA-Cre;R26R-LacZ;Ext1^e2fl/e2fl^* mice and E15.0 *Ext1^gt/gt^* embryos as well as the respective control littermates, embedded in Tissue-Tek O.C.T. medium (Sakura) and frozen at-56°C. AFM measurements were performed in the lab of Hauke Clausen-Schaumann at University of Applied Sciences, Munich, as published before (Prein, 2016) (Muschter, 2020). In brief, superficial, middle and deep zone of the AC were assessed on two separate sagittal tissue sections of each mouse with 1875 force-indentation curves in total distributed on 3 different 9μm² areas. A cantilever with a four-sided pyramidal tip geometry and 20 nm tip radius was used. A modified Hertz-Sneddon model was fitted onto the approach curve using Igor Pro (Version 6.3.7.2, WaveMetrics, Portland, USA) and the Young’s Modulus (YM) was extracted.

### Isolation and culture of MEFs

Murine embryonic fibroblasts (MEFs) were isolated from mouse embryos between E12.5 and E18.5. Head, limbs and internal organs were removed. The remaining tissue was homogenised and digested for 30Min at 37°C (0.3U/ml Collagenase NB4 (Serva), 5% FCS, 0.05% Trypsin-EDTA, in Dulbecco’s Phosphate Buffered Saline (DPBS)). The obtained cells were cultivated in DMEM:F12 1:1 (ThermoFisher Scientific) supplemented with 10% FCS (PAN) at 37°C and 5% CO_2_.

### Surfen treatment

To inhibit HS function, cells were treated with the small molecule antagonist *bis-2-methyl-4-amino-quinolyl-6-carbamide* (Surfen), supplied as Surfen hydrate (Sigma Aldrich). The number of H_2_O molecules in the hydrate is unknown, but it is determined by the production conditions and can thus be assumed to be stable.

For initial calculations we used the molecular weight of Surfen in its anhydrous state and made a stock solution of 0.37mg/ml Surfen hydrate in DMSO, corresponding to a maximum of 1mM Surfen. Our treatment conditions can thus be considered as ≤10, ≤5.0 and ≤2.5µM Surfen, containing 3.70, 1.85 and 0.925 µg/ml Surfen hydrate, respectively. For controls, we used the same volume of the solvent DMSO.

### Quantification of adherent cells

MEFs were seeded into culture medium at a density of 5×10^3^ cells/well in 24-well plates and cultured for 0.5, 1, 2, 16 or 24 hours (h). Cells were fixed with 4% PFA in DPBS for 10-15min at RT and stored covered with 1% PFA until processing. For nuclear staining, MEFs were c with 500ng/ml DAPI in ddH_2_O for 5min at RT and covered with DPBS for imaging using a reflected light fluorescence microscope (AxioObserver 7, Zeiss). Three images per well were acquired and the number of nuclei per field of view was quantified using CellProfiler 3.0.

### Life cell imaging

For the recording of life cell images, cells were seeded into Surfen- or DMSO-containing medium without phenol red at a plating density of 1.5×10^5^ cells per well on 8-well μ-slides (ibidi). After 0.5h pre-adhesion, acquisition was started and one image per minute was obtained by a phase contrast microscope (Ti Eclipse Epi, Nikon) at 5 positions per well for 1h from each condition.

### Immunofluorescence staining of MEFs

For immunofluorescence stainings, MEFs were seeded into 96-well plates suitable for fluorescence imaging (Corning 3614) at a density of 5×10^4^ of cells per well for a treatment duration of one hour or 2.5×10^4^ cells per well for a treatment duration of 24h. Cells were fixed with 4% PFA, permeabilised with 0.1% Triton X-100 in PBS and blocking was performed using 1% BSA in PBS. Primary antibodies against Paxillin (BD biosciences 610051) and YAP (Santa Cruz sc-101199) as well as appropriate secondary antibodies (Thermo Fisher, AlexaFluor) were used.

Nuclei were stained with 500ng/ml DAPI and Actin was labelled using Phalloidin-AlexaFluor633 (Thermo Fisher). Cells were post-fixed with 4%PFA and stored at 4°C covered with 1%PFA until imaging using a reflected light microscope (TIRF DualCam, Nikon).

### Biomimetic platforms

CHO *WT* (CHO-K1; (Puck, 1958)) and *pgsD-677* (Lidholt, 1992) cells were a gift from Kai Grobe. Biomimetic platforms were produced in Elisa Migliorini’s lab as published before (Sefkow-Werner, 2020). CHO cells were seeded at a density of 1×10^4^ cells per well of a biomimetic platform in 96-well plate format, cultured in DMEM:F12 (1:1) supplemented with 10% FBS and fixed using 4%PFA for 15min after 75min of adhesion. Staining was performed using a primary antibody against HS (clone F58-10E4, 370255-1 Amsbio) and an appropriate AlexaFluor-labelled secondary antibody (Thermo Fisher) as well as a fluorescence-labelled Phalloidin (Thermo Fisher).

### Preparation of primary murine chondrogenic progenitor cells and high-density culture

Limb buds of E10.5 – E11.5 C57Bl/6 mice were harvested, incubated with 1U/ml Dispase (Serva) for 15min at 37°C and dissociated (0.3U/ml Collagenase NB4 (Nordmark), 0.05% Trypsin-EDTA (Invitrogen), 5% FBS, in DPBS) for 30min at 37°C. The solution was passed through a cell strainer (70µm mesh width) and primary murine chondrogenic progenitor cells (pCh) resuspended at 2×10^7^ cells/ml. 2×10^5^ cells in 10µl were seeded per micromass into 24-well-plates. After 1h aggregation at 37°C, cells were cultured with 10% FBS, 1% Pen/strep (gibco), 1mM beta-glycerolphosphate and 0.25mM ascorbic acid phosphate in DMEM:F12 (1:1, Thermo Scientific) for chondrogenic differentiation (Underhill, 2014) (Schroeder, 2019).

On day 6, micromasses were fixed with 100% EtOH at-20°C for 20min and stained oN with Alcian Blue Staining Solution (TMS-010-C, Millipore). After washing, alcian blue was solubilised oN at RT using 6M guanide hydrochloride. The optical density of the resulting SN was measured at 595nm wavelength (GENios Pro plate reader, Tecan).

### Statistical Analysis

For statistical analyses, we used the probabilistic programming language Stan (version 2.32.2) together with R (version 4.4.2) (Team, 2025) (R Core Team, 2024) (Carpenter, 2017) to build experiment-specific models (Supp. File 01) and perform Bayesian inference to obtain posterior distributions of the model parameters. All models are fitted using four Markov chains with 2,000 iterations each, including 1,000 warm-up iterations. Model convergence was confirmed using the *R̂* diagnostic (*R̂* < 1.05), and model adequacy was assessed using posterior predictive checks. Posterior distributions of effect parameters 𝛿 are summarized by their median and 95% highest density intervals. To quantify evidence for differences between groups, we used a slightly adapted 𝜋-value (Kitanovski, 2020)

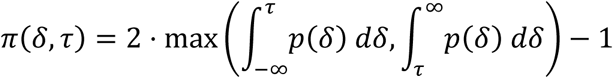

If 𝛿 represents the posterior distribution of a group difference, then 𝜋 = 1 indicates strong evidence for a true difference, whereas 𝜋 = 0 indicates no evidence for a difference. The output of the Bayesian analysis and number of analysed experimental replicates are listed in (Supp. File 02), means and standard deviation for plotting in (Supp. File 03).

## Results

### Altered composition of ECM synthezised by HS-deficient chondrocytes

In *Col2-rtTA-Cre;Ext1^e2fl/e2f^* mice (Jones, 2010) low Cre activity leads to the formation of clones of HS deficient chondrocytes surrounded by wild type-like AC (Fig. 1, A-D). The HS-deficient cells show an enlarged morphology and are surrounded by an increased GAG deposition as indicated by an intense Safranin O staining (Jones, 2010) (Severmann, 2020).

**Figure 1:**
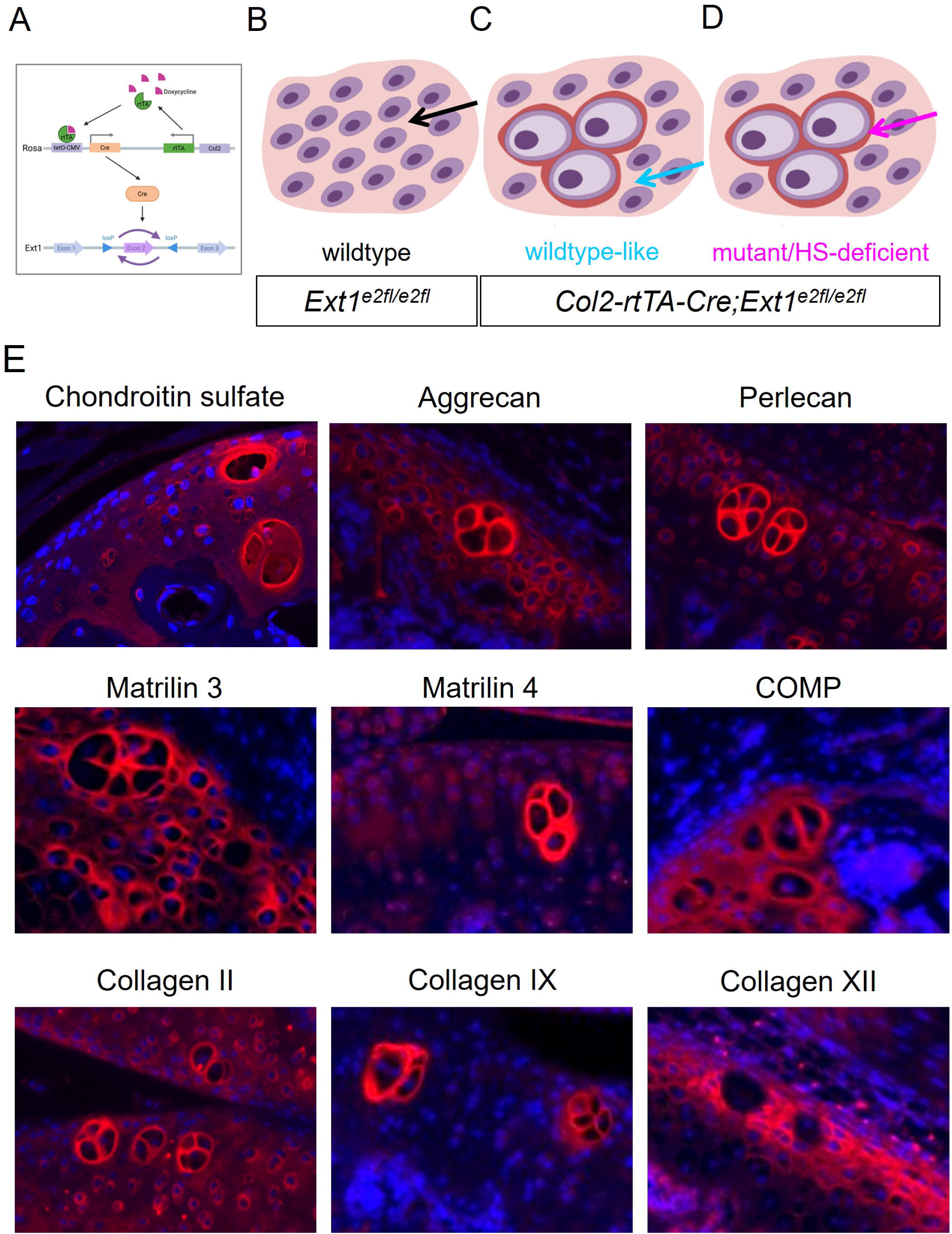
HS-deficient chondrocytes produce an ECM with markedly altered composition. (A) Schematic representation of clonal Ext1 deletion in transgenic *Col2-rtTA-Cre;Ext1^e2fl/e2fl^* mice (Buchholz, 2026a). (B, C, D) After induction of Cre activity at P8, clusters of enlarged, HS-deficient chondrocytes (D, magenta arrow) were detected in the AC of *Col2-rtTA-Cre;Ext1^e2fl/e2fl^* animals, but not *Ext1^e2fl/e2fl^* controls (B, black arrow). HS-deficient clusters were surrounded by wildtype-like tissue in the mutant mice (C, blue arrow). (E) Immunofluorescence stainings of sections of 4-week-old *Col2-rtTA-Cre;Ext1^e2fl/e2fl^* animals. Comparison of HS-deficient clusters to the surrounding wildtype-like tissue shows that levels of CS, Acan, Pcan, Matn-3, Matn-4, Collagen II and Collagen IX were increased in the mutant ECM, while Collagen XII was decreased and COMP remained unaltered.

**Supplementary Figure 1:**
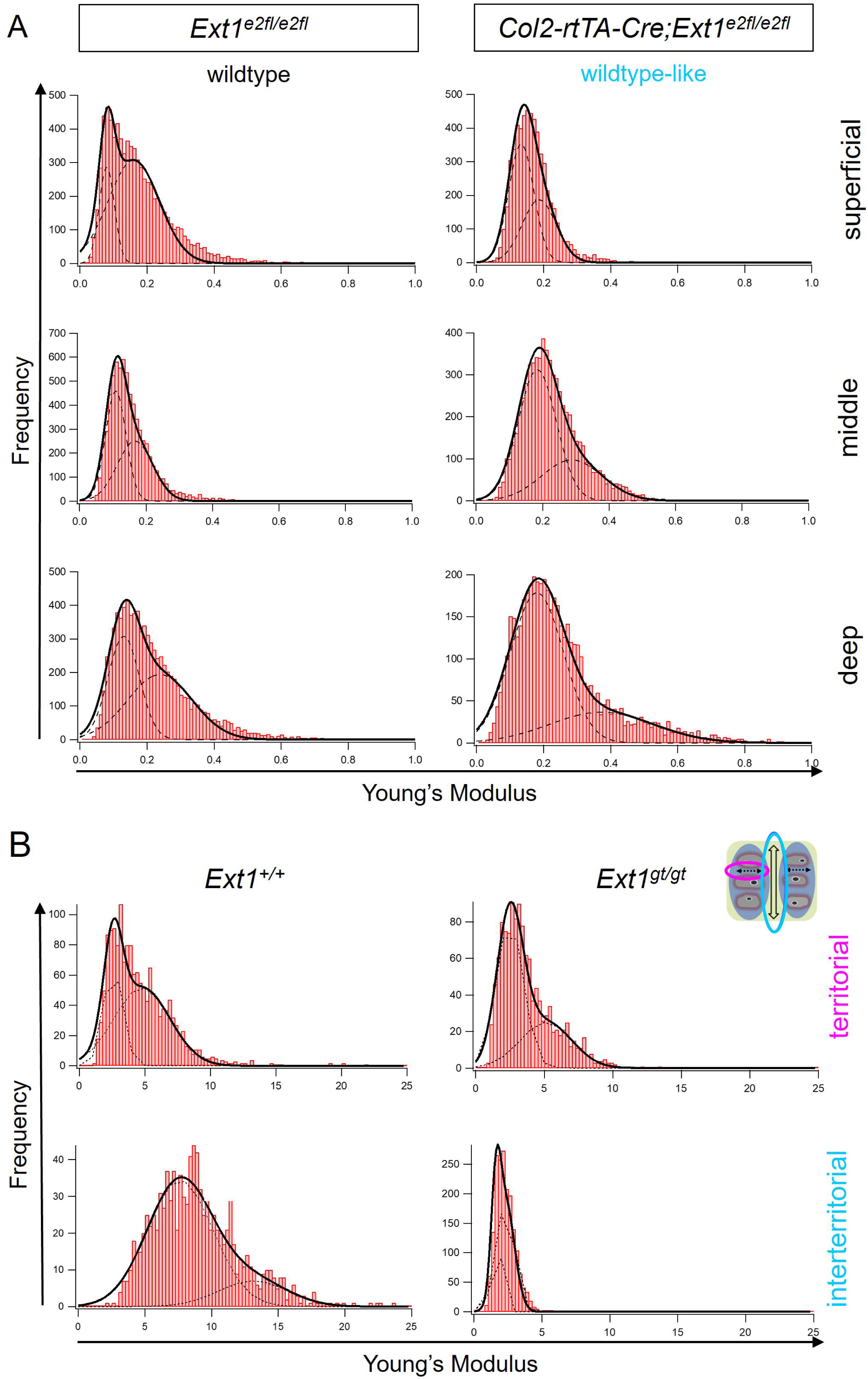
HS-deficient matrix of an *Ext1^gt/gt^* embryo shows decreased Young’s Modulus (YM) Histograms show YM detected by AFM analysis of native, unfixed tissues. The YM was measured in three 9µm² areas per mouse on two distinct sagittal sections, resulting in 1875 data points per individuum. (A) Superficial, middle and deep zone of the AC of 4-week-old *Col2-rtTA-Cre;Ext1^e2fl/e2fl^*mice and wildtype littermates (*Ext1^e2fl/e2fl^*) were analysed. The wildtype-like matrix of the mutant mice showed a slightly higher YM compared to wildtype mice. It was not possible to measure the YM of the HS-deficient ECM in these samples. n=3. (B) Territorial and interterritorial matrix of the GP of one *Ext1^+/+^* and one *Ext1^gt/gt^* embryo were assessed at E15.0. While the stiffness of the territorial matrix seemed similar between the two genotypes, the interterritorial ECM was markedly softer in the *Ext1^gt/gt^*mutant. n=1.

To investigate the composition of the HS-deficient ECM in more detail, we analysed the levels of typical cartilage ECM components CS, Acan, Pcan, Matn-3, Matn-4, Col2, Col9, Col12 and COMP on sagittal sections of knees of 4w old *Col2-rtTA-Cre;Ext1^e2fl/e2fl^* mice. Due to the low Cre activity, HS-deficient and wildtype-like cells can be compared on the same section, bypassing slide-to-slide variations of fluorescence intensities (Fig. 1, B-D). All components were detected in the ECM surrounding wildtype-like chondrocytes. In line with the increased Safranin O staining, the level of CS and of the GAG-carrying proteins Acan and Pcan were elevated in the ECM surrounding the HS-deficient clones. Interestingly, besides the GAG-carrying proteins, structural components of the Collagen network, including Col2, Col9 and the Collagen-associated proteins Matn-3 and Matn-4, were also increased. In contrast, COMP was detected at similar levels in HS-deficient and HS-containing matrix, while Col12 seemed to be absent in the ECM of mutant cells (Fig. 1, E). Taken together, these findings show that HS-deficient chondrocytes synthesize a matrix of markedly altered composition.

### Reduced matrix stiffness of HS-deficient cartilage matrix

To investigate how the aberrant composition affects the mechanical properties of the mutant ECM, we analysed the stiffness of the joint cartilage in fresh-frozen samples of 4w old mice by AFM. Separate measurements were performed in the superficial, middle and deep zones of the AC. In *Ext1^e2fl/e2fl^* control animals (Fig. 1B), the bimodal distribution of the YM increased from the superficial zone to the deep zone. AFM analysis of the wildtype-like chondrocytes surrounding mutant cells of *Col2-rtTA-Cre;Ext1^e2fl/e2fl^*mutants (Fig. 1C) showed an increased stiffness in the three zones compared to control littermates (Fig.1B; Fig. S1A). Interestingly, the matrix directly surrounding the mutant cells (Fig. 1D) was too soft to be measured in the native samples.

To overcome this limitation, sections of fresh frozen samples were fixed with 4% PFA, resulting in a severely increased matrix stiffness in control and mutant animals compared to the fresh-frozen tissue. (Fig. S1A, Fig. 2A). The matrix surrounding wildtype-like cells of *Col2-rtTA-Cre;Ext1^e2fl/e2fl^* animals was still slightly stiffer than that of control littermates but the difference in stiffness observed in fresh-frozen samples seemed to be masked to some extent by the fixation procedure (Fig. 2A). Importantly, the ECM surrounding the HS-deficient cells could be measured after fixation and showed a considerably decreased YM compared to each of the AC zones (Fig. 2B). Similarly, analysis of an E15.5 *Ext1^gt/gt^* embryo (Mitchell, 2001) (Koziel, 2004), which produce about 20% of CS in chondrocytes (Yamada, 2004;Bachvarova, 2020) also revealed a markedly reduced matrix stiffness in the cartilage anlage compared to an *Ext^+/+^* littermate. While the territorial matrix (highlighted in magenta) exhibited a similar stiffness as that of the wildtype control, a clearly reduced YM was detected in the interterritorial matrix (highlighted in blue), supporting a role of HS in regulating ECM composition in chondrocytes (Fig. S1B).

**Figure 2:**
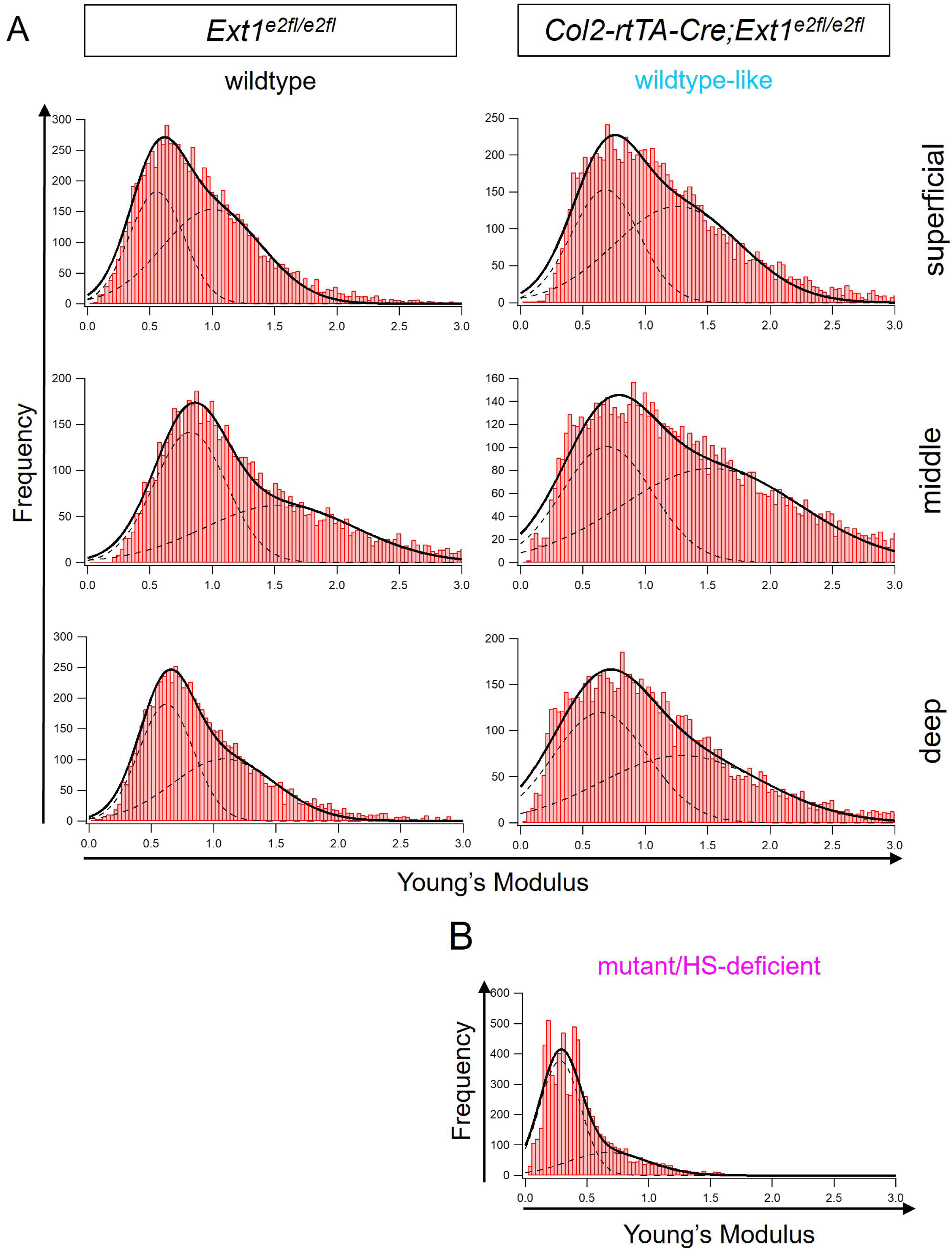
HS-deficient matrix of *Col2-rtTA-Cre;Ext1^e2fl/e2fl^* mice is soft. Histograms show YM detected by AFM analysis of PFA-fixed tissue, which resulted in an overall increase of tissue stiffness compared to native tissues. The YM was measured in three 9µm² areas per mouse on two distinct sagittal sections, resulting in 1875 data points per individuum. (A) Superficial, middle and deep zone of the AC of 4-week-old *Col2-rtTA-Cre;Ext1^e2fl/e2fl^* mice and *Ext1^e2fl/e2fl^*littermates were analysed. The wildtype-like matrix of *Col2-rtTA-Cre;Ext1^e2fl/e2fl^*animals showed a slightly higher YM than the ECM of wildtype littermates. (B) The HS-deficient matrix of the *Col2-rtTA-Cre;Ext1^e2fl/e2fl^* mice was markedly softer compared to the surrounding wildtype-like ECM and to wildtype tissue.

### Increased levels of Integrin pathway components in HS-deficient chondrocytes

Our results show that the loss of HS in chondrocytes is compensated by an upregulation of distinct ECM components. This leads to the question how chondrocytes sense the composition of the surrounding matrix.

Integrin dimers are major sensors of ECM quality and composition. Therefore, we analysed protein levels of the Integrin pathway components ItgB1, Src, Erk, pErk, FAK and pFAK, and NMII, which is part of contractile SF anchored to FA, by immunofluorescence on sagittal sections of 4w old *Col2-rtTA-Cre;Ext1^e2fl/e2fl^* mice.

ItgB1, the most prominent β subunit found in chondrocytes (Salter, 1992; Woods, 1994), was detected at similar levels in wild type and HS-deficient articular chondrocytes. Notably, ItgB1 was clustered along the cell membrane in a punctate fashion, pointing to an Integrin-dependent formation of FA-like structures in chondrocytes, *in vivo* (Fig. 3A, examples highlighted by arrowheads). In contrast, FAK and its activated, phosphorylated form (pFAK), Src kinase, ERK, pERK, and NMII showed homogenously elevated levels in HS-deficient clones compared to the surrounding wild type cells (Fig. 3 A, B) indicating elevated signalling activity of integrin related pathways.

**Figure 3:**
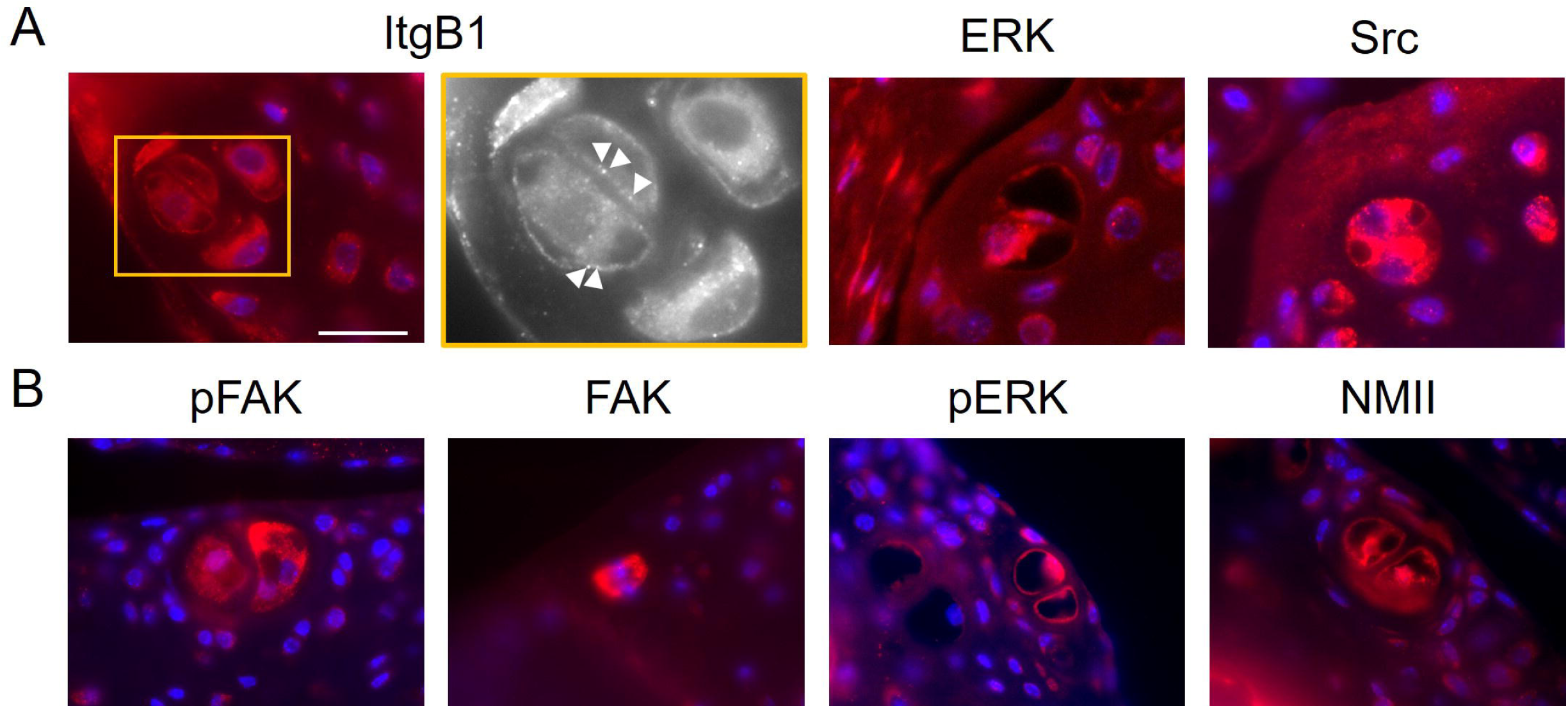
Levels of Integrin pathway components are increased in HS-deficient ECM. Immunofluorescence stainings of Integrin pathway components on sections of *Col2-rtTA-Cre;Ext1^e2fl/e2fl^* mice. (A) The level of ItgB1 was similar between wildtype-like tissue and HS-deficient clusters of 4-week-old *Col2-rtTA-Cre;Ext1^e2fl/e2fl^* animals and ItgB1 was observed in a punctate distribution along the cell membrane. Increased levels of ERK and Src were detected in HS-deficient clones. Scale bar: 20µm. (B) Increased levels of pFAK, FAK, pERK and NMII were detected in HS-deficient clones of 8-week-old *Col2-rtTA-Cre;Ext1^e2fl/e2fl^* mice.

### Impaired cell adhesion and migration upon inhibition of HS function

To further analyse the influence of HS on Integrin-dependent processes, we switched to an *in vitro* model using primary MEFs, a well described system to study integrin-dependent cell-substrate adhesion, spreading and migration. We mimicked the loss of HS by treating the cells with the small molecule antagonist Surfen (Schuksz, 2008). To investigate the effect of Surfen on adhesion, MEFs were seeded into medium containing ≤10, ≤5.0 and ≤2.5µM Surfen or control medium (control), and the number of adherent cells was quantified after 0.5, 1, 2, 16 and 24h using the image analysis software CellProfiler (Stirling, 2021). In both, control and Surfen-treated cultures, the numbers of cells adhering to the surface of the culture dish increased over time. However, compared to the control the numbers of adherent cells were reduced in Surfen containing medium in a concentration dependent manner. The effect was strongest at early timepoints 0.5 and 1h. It became less pronounced at 2h and was largely compensated after 16 and 24h, supporting a role of HS in the early adhesion process (Fig. 4A, B). The lowest concentration with a clear effect on early cell adhesion (0.5 to 2h), ≤5µM, was used for all further experiments.

**Figure 4:**
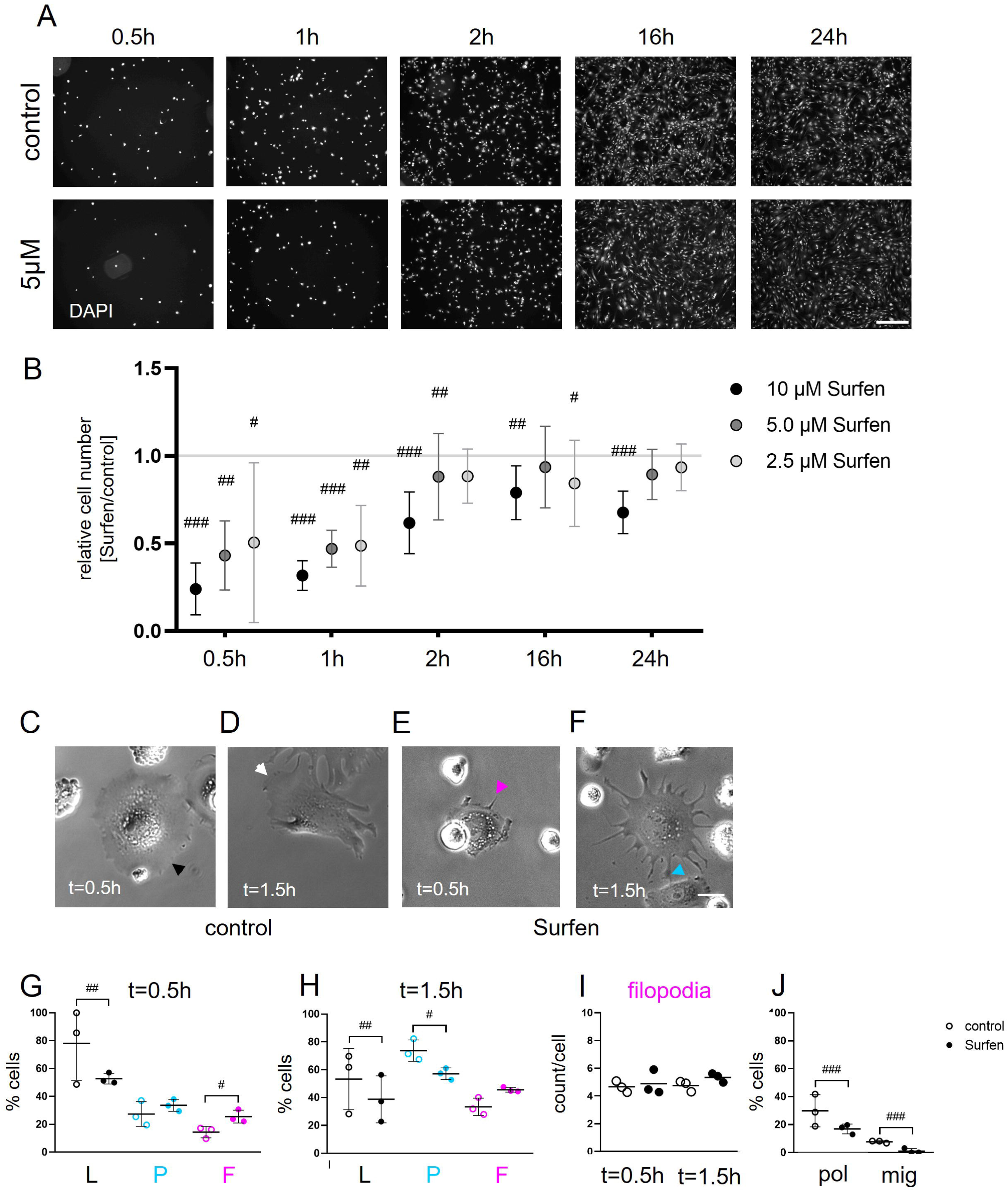
Inhibition of HS-function impairs Integrin-dependent processes. Analysis of MEFs treated with the HS-antagonist Surfen. (A, B) Investigation of the number of cells adhering to the cell culture dish upon Surfen-treatment. (A) Detection of adherent cells via nuclear staining at 0.5, 1, 2, 16 and 24h post seeding. The numbers of adhering MEFs increased over time in control and Surfen-treated samples. Scale bar: 200µm. (B) Quantification of adherent cells upon treatment with ≤10, ≤5 and ≤2.5µM Surfen relative to the control. During early cell adhesion up to 2h post seeding, Surfen reduced cell-substrate adhesion in a dose-dependent manner. This effect was largely compensated at later time points. n = 2. Mean with SD displayed. (C-J) Life cell imaging of MEFs treated with ≤5µM Surfen from 0.5 to 1.5h post seeding. (C-F) Phase contrast images from control and Surfen-treated MEFs at 0.5 and at 1.5h post seeding. Distinct membrane morphologies were observed: Smooth, lamellipodium-like cell seems (C, black arrowhead; D, white arrowhead), thin, filopodia-like protrusions (E, magenta arrowhead) and branched, pseudopodia-like protrusions (F, blue arrowhead). Scale bar: 20µm. (G, H) Quantification of the number of cells showing lamellipodia- (L), pseudopodia- (P) and filopodia-like (F) protrusions upon treatment with Surfen relative to control medium. The proportion of cells with lamellipodia-like protrusions decreased, while the percentage of MEFs with filopodia-like protrusions increased in Surfen-treated samples at 0.5 and at 1.5h post seeding. The percentage of cells with pseudopodia-like protrusions showed no consistent changes, it was slightly increased at 0.5 and decreased at 1.5h post seeding. (I) Quantification of the number of filopodia on each filopodia-forming cell showed no difference between treatment and control at both time points. (J) Quantification of polarising and migrating MEFs during time lapse from 0.5 to 1.5h post seeding. In Surfen-treated samples, proportions of polarising and migrating cells were reduced. Black lines indicate means with SD. n = 3. # π ≥ 0.90, ## π ≥ 0.95, ### π = 1.

Cell spreading and migration rely on integrin-dependent substrate adhesion and reorganization of the cytoskeleton, leading to changes in cell shape and membrane morphology (Vicente-Manzanares, 2009) (Parsons, 2010). We quantified the formation of membrane protrusions upon Surfen treatment by life cell imaging of the critical adhesion period from 0.5h to 1.5h post seeding. Cell phenotypes were defined depending on membrane morphologies: Broad cell protrusions with a smooth edge were termed “lamellipodia-like” (Fig. 4C, D; black and white arrowheads), thin finger-shaped protrusions “filopodia-like” (Fig. 4E; magenta arrowhead) and broader, branched protrusions “pseudopodia-like” (Fig. 4F; blue arrowhead). Quantification at 0.5 and at 1.5h after seeding showed that the proportion of cells with a lamellipodia-like membrane phenotype was decreased upon Surfen treatment compared the control samples. Surfen treatment did not result in consistent effects on the proportion of cells forming pseudopodia-like protrusions, which showed a tendency to be increased at 0.5h while being decreased at 1.5h. In contrast, MEFs with filopodia were increased at 0.5h and still showed a similar tendency at 1.5h (Fig. 4G, H). Since the number of cells generating filopodia-like protrusions seemed to be increased by treatment with Surfen at both time points, we quantified their number on each filopodia-carrying cell but did not find a difference to control cultures (Fig. 4I). Surfen does thus not affect the number of filopodia-like protrusions per cell but specifically the number of cells forming such protrusions, indicating a delay on the transition between initial cell adhesion and spreading to cell polarisation and finally migration. Therefore, we followed individual cells throughout the whole time interval and revealed less polarising and migrating cells in the Surfen treated cultures (Fig. 4J).

Taken together, these data reinforce the hypothesis that HS is essential for Integrin-dependent cell adhesion and the initiation of migration.

### Endogenous HS necessary for cell polarization

The positively charged Surfen inhibits HS function based on electrostatic interaction with its negatively charged sulfate groups (Schuksz, 2008). To rule out unspecific effects of Surfen on other negatively charged structures, we analysed the polarisation of CHO *pgsD-677* cells, which do not produce HS (Lidholt, 1992), and compared it to CHO *wildtype (WT)* cells. The lack of HS in CHO *pgsD-677* cells was confirmed by immunofluorescence with the HS-specific antibody 10E4. CHO *WT* cells showed a clear HS staining, while no signal was detected for CHO *pgsD-677* cells (Fig.5A, B). We seeded the cells onto biomimetic platforms functionalised with a defined surface concentration of 10ng/cm^2^ RGD peptide, to facilitate cell adhesion in a standardized manner (Fig. 5C) (Sefkow-Werner, 2020) and visualised the actin cytoskeleton with fluorescence-conjugated Phalloidin after 75min of adhesion. While the *WT* cells showed a polygonal morphology, CHO *pgsD-677* cells appeared rather round (Fig. 5D, E). To quantify the observed difference in cell shape, we measured the cell ellipticity, defined as the short axis of the cell divided by the long axis, using ImageJ (Schneider, 2012). A ratio close to 1 indicates a round cell morphology, while a lower ratio can be interpreted as a polarised, more elongated cell shape. We found that CHO *WT* cells had a more elliptic shape, while CHO *pgsD-677* cells were less elliptic. To test if exogenous HS can rescue the altered shape of HS-deficient cells, CHO *WT* and *pgsD-677* cells were plated on modified platforms presenting RGD peptides together with HS at a surface concentration of 14.4 ±0.5ng/cm^2^ (Fig. 5C). On this surface, CHO *WT* cells also lost their elliptic shape (Fig. 5D, H), likely due to the increased negative charge of the surface. Importantly, CHO *pgsD-677* mutants developed a similarly roundish shape as on the RGD surface (Fig. 5G, H) strongly indicating that external HS is not sufficient to compensate for the lack of cellular HS.

**Figure 5:**
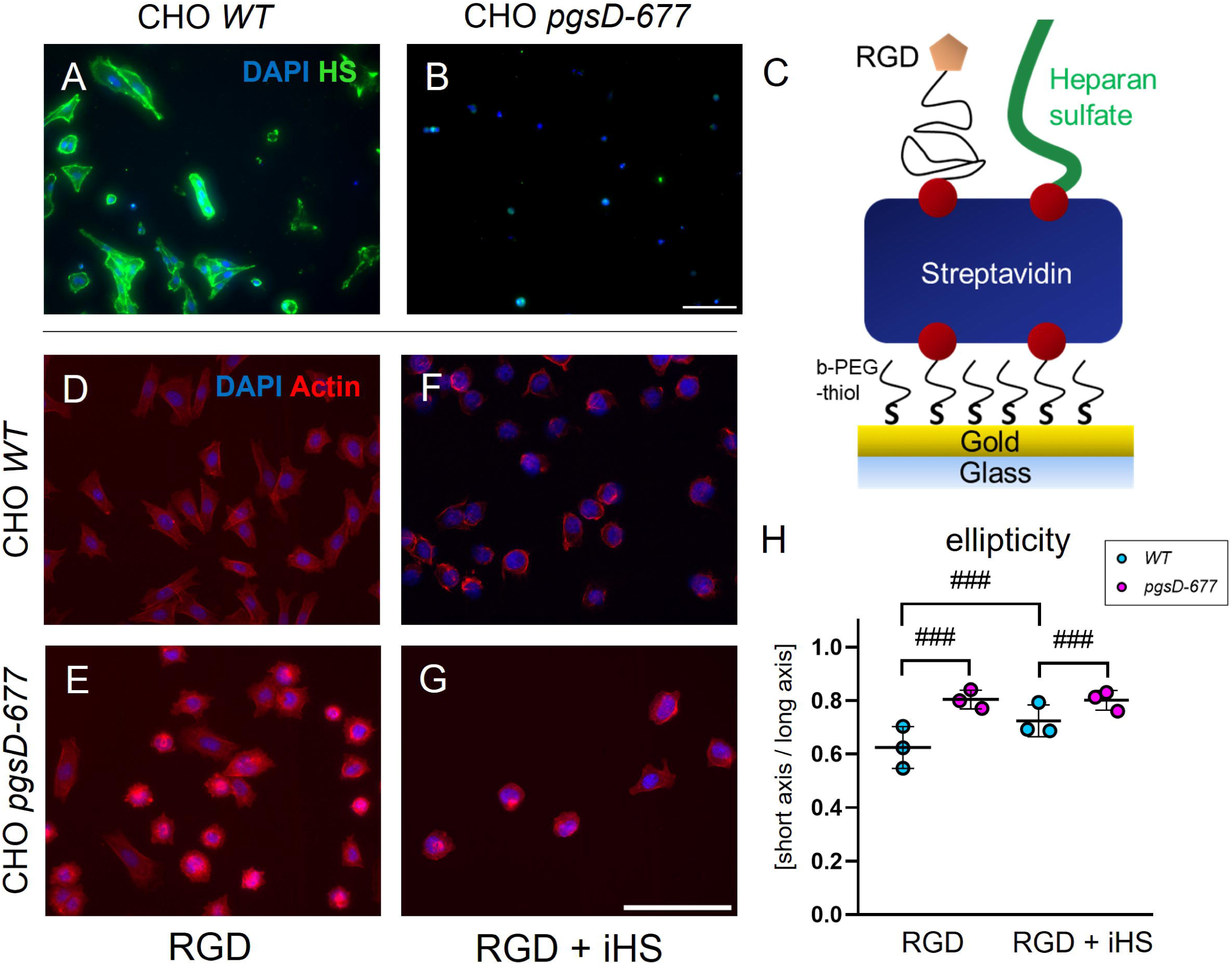
Loss of endogenous HS-synthesis cannot be rescued by exogenous presentation of HS. Fluorescence images of CHO wildtype and HS-deficient CHO *pgsD-677* cells 1.25h post seeding. (A, B) Detection of HS by immunofluorescence staining with anti-HS-antibody clone 10E4. HS can be detected on CHO WT but not CHO *pgsD-677* cells. Scale bar: 100µm. (C) Schematic representation of biomimetic platforms presenting RGD peptides and immobilised HS (iHS) (D-F) Visualisation of the actin cytoskeleton using a fluorescence-labelled Phalloidin. On RGD, CHO WT cells show an elongated phenotype compared to HS-deficient CHO *pgsD-677* cells. When RGD and iHS are presented, both CHO WT and CHO *pgsD-677* cells display a rounded morphology. Scale bar: 100µm. (H) Quantification of ellipticity defined as short cell axis / long cell axis. On RGD-presenting platforms, CHO WT cells are more elliptic with a lower mean value, and CHO *pgsD-677* cells are less elliptic with a higher mean. On platforms with RGD and iHS, CHO *pgsD-677* cells show a similar rounded phenotype compared to RGD only, while CHO WT cells become less elliptic. Black lines indicate means with SD. n = 3: # π ≥ 0.90, ## π ≥ 0.95, ### π = 1.

### HS required for formation of focal adhesions and stress fibres during early cell adhesion and spreading

Cell polarisation and migration require a stable adhesion to the underlying substrate and the reorganisation of the cytoskeleton to generate traction within the cell. These processes rely on the Integrin-dependent formation of focal adhesions (FAs) at the cell-substrate interface and the assembly of contractile stress fibres (SFs) (Vicente-Manzanares, 2009) (Parsons, 2010). To detect the presence of FA, MEFs were seeded into control or Surfen containing medium and analysed after different time periods by immunofluorescence. FAs were detected using an antibody against Paxillin, an adaptor protein of the FA complex (Schaller, 2001).

In control cells, Paxillin-positive FA were clustered at the cell periphery (Fig. 6A, blue arrowhead) while in Surfen-treated cells the Paxillin signal was more homogenously distributed throughout the cell body (Fig. 6A, magenta arrowhead). Quantification of cells with at least one FA revealed that the number of cells forming FAs was reduced in the presence of Surfen compared to control cells at 1h after seeding. At 24h, no clear effect was observed, and the number of cells with FA was similar in treated and untreated cells (Fig. 6B).

**Figure 6:**
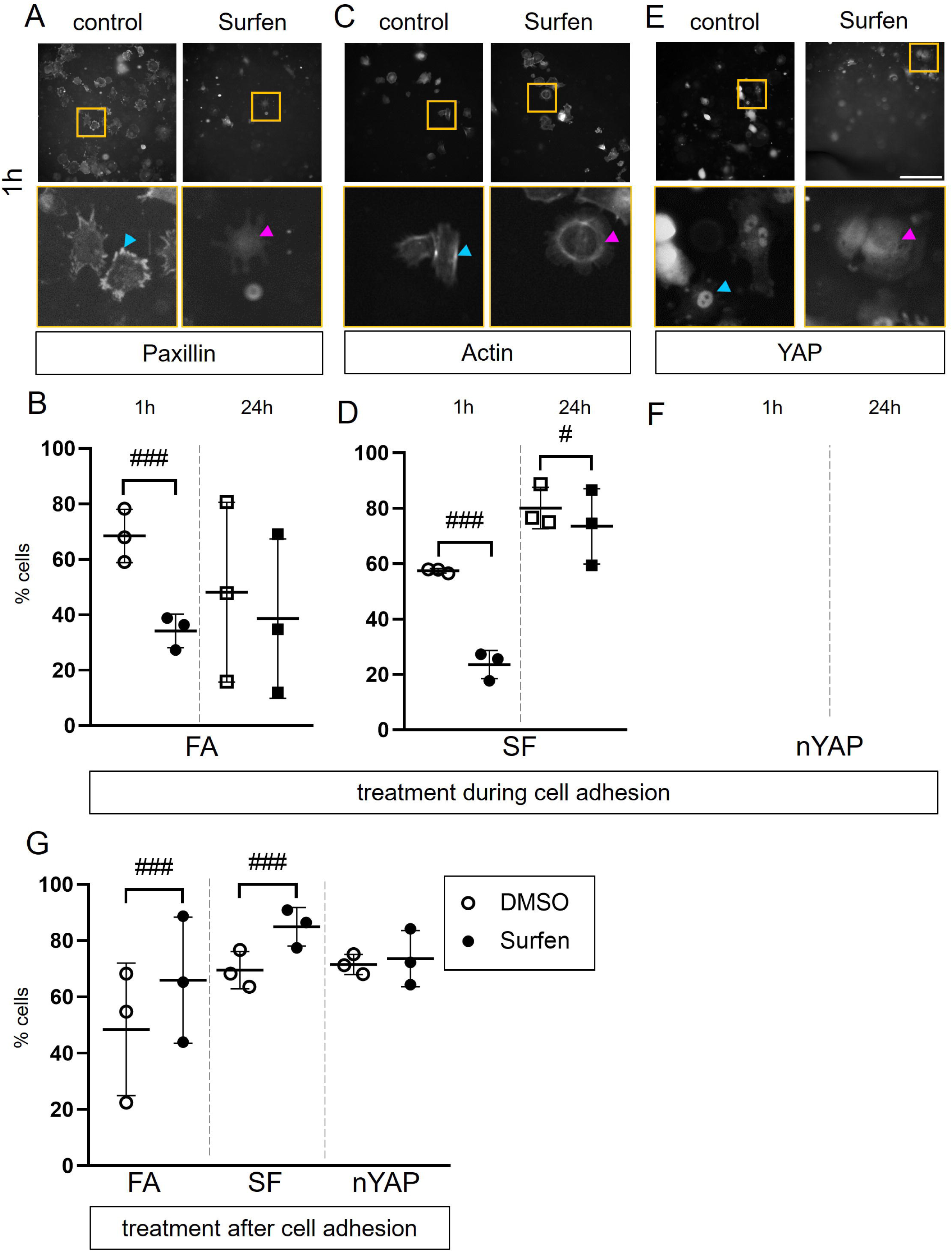
HS inhibition during and after cell adhesion induces opposing effects. Investigation of the effect of HS inhibition on the formation of FAs, SFs and nuclear localisation of YAP (nYAP). (A, C, E) Fluorescence images of MEFs seeded into medium with ≤5µM Surfen or control medium. FA were detected by staining against Paxillin, the actin cytoskeleton visualised using a fluorescence-labelled Phalloidin and YAP was stained by immunofluorescence. Scale bar: 200µm. (B, D, E) Quantification of the numbers of MEFs seeded into Surfen-containing medium forming FA, SF and showing a clear nuclear signal for YAP. At 1h post seeding FA, SF and nYAP were reduced in the Surfen treated samples. After 24h, there was no effect on FA and the proportion of cells forming SF was largely compensated while the proportion of cells showing nYAP remained markedly decreased. (G) When already adherent cells were treated with Surfen, quantification showed increased percentages of MEFS with FA and SF, while there was no effect on nYAP. (B, D, E, F) n = 3. Black lines indicate means with SD. # π ≥ 0.90, ## π ≥ 0.95, ### π = 1.

Next, we quantified SF formation using a fluorescence-conjugated Phalloidin to visualise actin (Verderame, 1980). In many control cells, we found prominent linear SF along the primary cell axis (Fig. 6C, blue arrowhead), while in Surfen treated samples cells more often showed disorganized actin filaments in the cell periphery (Fig. 6C, magenta arrowhead). We quantified the number of cells with prominent SF along the primary axis of the cell body. In general, compared to the earlier time point of 1h adhesion, a higher proportion of cells formed SF after 24h of adhesion, underlining the transition of MEFs from initial cell adhesion to migration and polarisation. Compared to controls, the formation of SF was reduced in Surfen-treated MEFs after 1h of adhesion. This the effect was mostly compensated after 24h but still statistically clear (Fig. 6D). Taken together, treatment of MEFs with Surfen results in a reduced formation of FA and SF during cell adhesion. This is in line with the observed inhibition of cell adhesion, polarisation and migration described above (see Fig. 4).

### Subcellular localisation of YAP controlled by HS function

Cell adhesion and the formation of SFs are directly linked to gene regulation via the transcriptional activator YAP. As Surfen treatment led to the changes in SF formation (Fig. 6C, D), we quantified the number of cells with a clear nuclear localisation of YAP detected by antibody staining in control and Surfen-treated cells. We observed cells with a strong nuclear signal (Fig. 6E, blue arrowhead) while others displayed a uniform distribution of YAP within the entire cell body (Fig. 6E, magenta arrowhead).

The percentage of MEFs with a nuclear localisation of YAP was similar under control conditions at 1h and at 24h after seeding. At both time points, it was reduced in the presence of Surfen, indicating that YAP localisation relies on HS function during early cell adhesion and that the effect is persistent even after the cells fully adhered.

### Distinct effects of HS during initial cell adhesion and at later stages of attachment

While the formation of FA and SF in MEFs were impaired at early time points *in vitro*, HS-deficient cells of *Col2-rtTA-Cre;Ext1^e2fl/e2fl^* mice showed elevated levels of Integrin pathway components, including FAK, pFAK and NMII, *in vivo* (see Fig. 3). To analyse the effect of an antagonization of HS-function after completion of the adhesion process, we allowed cells to adhere for 24 h before adding Surfen to the culture for 1h. Interestingly, the formation of FA and SF was increased in the Surfen-treated samples while YAP localization was not affected (Fig. 6G). Taken together these effects show that loss of HS-function results in distinct effects during early cell adhesion and in less dynamic adherent cell states.

### YAP signalling regulates GAG production in chondrocytes

Our results from treatment of MEFs with Surfen *in vitro* showed distinct effects of HS function on YAP localisation during cell adhesion and in already adherent cells. This led us to investigate the level of YAP in clusters of HS-deficient chondrocytes *in vivo.* We detected YAP on sections of 4w old *Col2-rtTA-Cre;Ext1^e2fl/e2fl^* mice by IF and found an increased level of YAP in HS-deficient chondrocytes in the AC (Fig. 7A).

**Figure 7:**
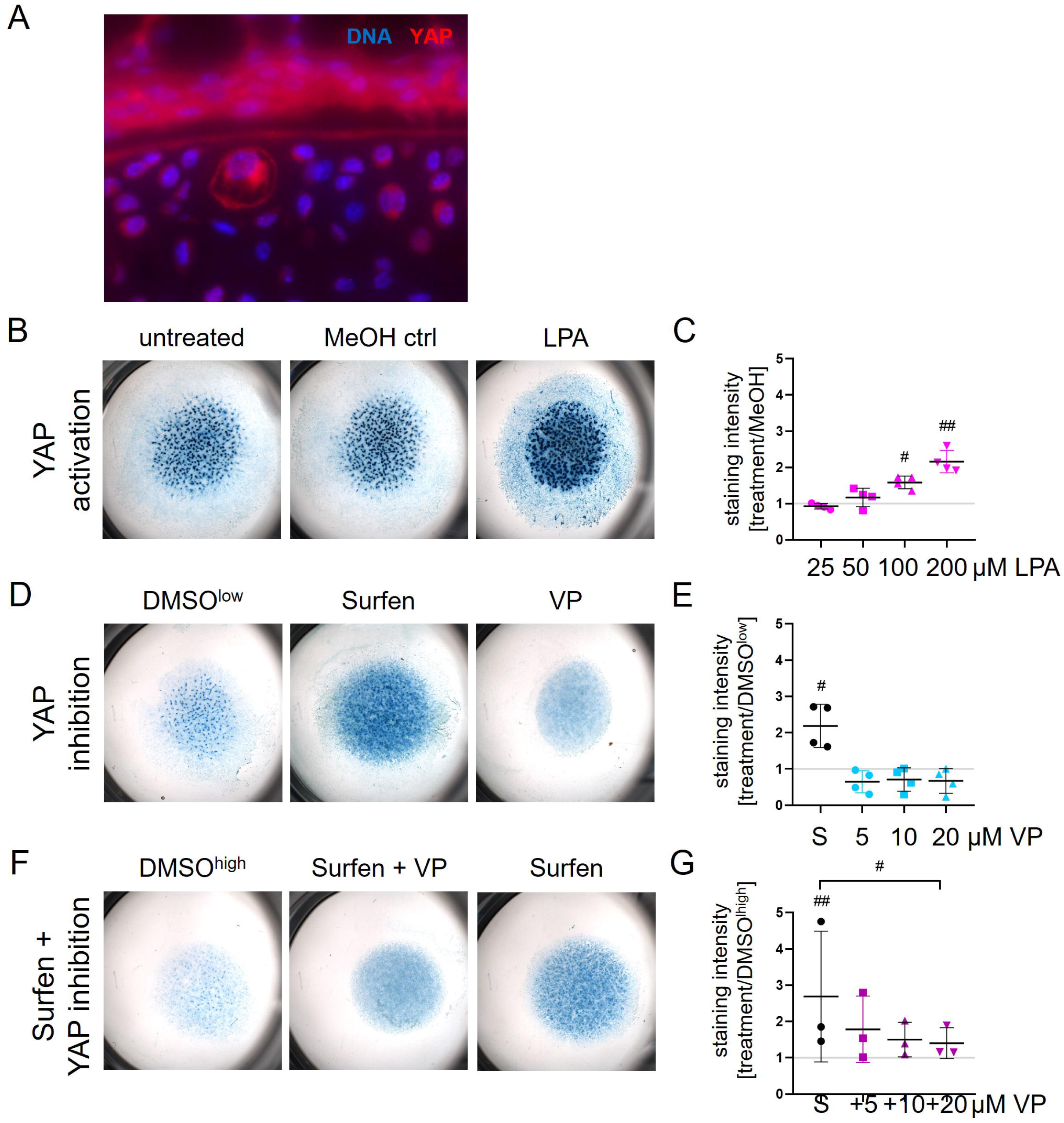
YAP controls GAG synthesis in chondrogenic cells. (A) Immunofluorescence staining of YAP on sections of 4-week-old *Col2-rtTA-Cre;Ext1^e2fl/e2fl^* mice showed an elevated level of YAP in HS-deficient clusters. (B, D, F) Alcian blue staining of GAGs in high density micromass cultures from primary murine chondroprogenitors (pCh). (C, E, G) Quantification of solubilized Alcian blue relative to controls. (C) Treatment with 25, 50, 100 and 200µM of the YAP activator LPA induced GAG production in a dose-dependent manner compared to the MeOH control. (E) Similarly, ≤7.5µM Surfen resulted in increased GAG synthesis. Treatment with 5, 10, 20µM of the YAP inhibitor VP slightly reduced GAG deposition compared to the solvent control. (G) When micromasses are treated with Surfen and VP simultaneously, VP blocks the effect of Surfen in a dose-dependent manner. (C, E, G) n = 3-4. Black lines indicate means with SD. # π ≥ 0.90, ## π ≥ 0.95, ### π = 1.

We showed that reduced HS function affects the nuclear localisation of YAP, while it has been reported that inhibition of HS leads to an upregulation of GAG production in chondrogenic cells. We thus asked if YAP mediates the observed upregulation of GAGs downstream of HS. To analyse this, we treated micromass cultures of primary murine chondroprogenitors (pCh) with Surfen and pharmacological modulators of YAP signalling and quantified the relative GAG amount by Alcian blue staining. As expected, Surfen treatment increased the GAG production compared to controls (Huegel, 2013b). Similarly, treatment with 25, 50, 100 and 200µM of the YAP-activator lysophosphatidic acid (LPA; (Yu, 2012) increased GAG synthesis in a dose-dependent manor with a statistically clear increase at concentrations of 100 and 200µM (Fig. 7B, C). In contrast, the addition of 5, 10 and 20µM of the YAP-inhibitor Verteporfin (VP; (Wang, 2016)) led to a mild but not statistically evident decrease of the GAG amount at all tested concentrations (Fig. 7 D, E). Still, combined treatment with Surfen and VP abolished the effect of Surfen on GAG accumulation in a dose-dependent manner with a statistically clear difference to Surfen treatment alone at 20µM VP (Fig. 7F, G). This finding indicates that YAP acts downstream of HS-function and contributes to the upregulation of GAG production in HS-deficient chondrocytes.

## Discussion

Clonal loss of HS synthesis in chondrocytes leads to the accumulations of an PG-rich, soft ECM in *Col2-rtTA-Cre;Ext1^e2fl/e2f^* mice. We investigated how the cells sense the lack of HS and how this is translated into increased ECM production. *In vitro*, analyses of cell adhesion, migration and the formation of FA and SF in MEFs indicated impaired Integrin-dependent processes after inhibiting HS function with Surfen and a lack of functional HS on the cell surface could not be rescued by presentation of exogenous HS. Interestingly, treatment of adherent cells with Surfen increased FA and SF formation, in accordance with the accumulation of Integrin pathway components *in vivo*. Furthermore, we demonstrated that the subcellular localisation of YAP depends on HS-function and that the Surfen induced GAG production can be antagonized by inhibition of YAP, showing that YAP is not only a downstream effector of HS function but directly controls GAG synthesis.

### HS affects Integrin-dependent processes in distinct ways during early cell adhesion and in mature tissue

Investigation of *Col2-rtTA-Cre;Ext1^e2fl/e2f^* mutants demonstrated an accumulation of Integrin pathway components in HS-deficient clones *in vivo*. To further investigate the role of HS for Integrin function, we treated MEFs with the HS-antagonist Surfen *in vitro*. During the adhesion process this resulted in reduced numbers of adhering, polarising and migrating MEFs and an impaired formation of FA and SF. Likewise, it has been reported that endothelial cells show reduced numbers of SF and decreased FA sizes after heparinase II treatment (Moon, 2005). While the direct role of HS is rarely investigated, many studies demonstrated the importance of the HS-carrying core protein family of Syndecans (Sdcs) for Integrin signalling (Pap, 2013). Sdc4 is a component of FAs, co-localising with ItgB1 (Oh, 2004) (Sarrazin, 2011) and fibroblasts lacking Scd4 show a reduced formation of FA and SF (Gopal, 2010). Accordingly, overexpression of Sdc4 in CHO cells induced an increased formation of FA and SF (Longley, 1999), confirming the essential role of HSPGs for Integrin-dependent cell adhesion.

We found that loss of endogenous HS in CHO *psgD-677* cells was not rescued by the presentation of HS on the adhesion surface, highlighting that cell-surface HS are essential for cell adhesion, spreading and polarisation. This is in line with our observations *in vivo*, as the lack of HS in mutant clusters is not compensated by the surrounding wildtype-like cells in *Col2-rtTA-Cre;Ext1^e2fl/e2fl^* mice. HSPGs, especially such with Sdc core proteins, have been shown to act as co-receptors for Integrin signalling, Integrin-dependent cell migration and downstream reorganisation of the cytoskeleton (Hassan, 2021). For instance, Sdc-1 interacts with α6β4 integrin dimers enhancing survival and motility (Wang, 2014) (Wang, 2015) and Sdc-1 also interacts with αvβ3 integrins, inducing migration (Pasqualon, 2015) in tumour cells. Similarly, Sdc-4 promotes mobility (Wang, 2014) (Wang, 2015) and complex formation of Sdc-2 and α5β1 induces formation of SFs (Munesue, 2002) in cancer cells. Moreover, the intracellular domains of Sdcs interact with downstream factors of Integrins. Sdc-4 induces signalling through RhoA and Rac1 GTPases, leading to the formation of FAs (Dovas, 2010), and enhancing migration (Bass, 2007).

An interplay between HS and Integrin is also highlighted by the similar phenotypes of ItgB1- and HS-deficient mouse mutants. We previously reported a disorganised columnar zone in the GP of *Ext1^gt/gt^* mutants (Mitchell, 2001), which fail to form longitudinal septae of interterritorial matrix (Koziel, 2004), resembling mice with a chondrocyte-specific deletion of ItgB1 (*Col2-Cre;β1^fl/fl^*) (Aszodi, 2003). This is reflected by our analysis of an E15.0 *Ext1^gt/gt^* embryo and its *Ext1^+/+^* littermate: the interterritorial matrix between columns of proliferating chondrocytes in the GP is typically stiffer than territorial (Prein, 2016), a characteristic that was lost in the *Ext1^gt/gt^* mutant where the stiffness of the interterritorial matrix was clearly reduced. In postnatal samples, the YM increased with the depth of the AC as expected (Muschter, 2020), in both *Col2-rtTA-Cre;Ext1^e2fl/e2fl^* mice and *Ext1^e2fl/e2fl^* littermates. The wildtype-like matrix of the mutants was slightly stiffer than the ECM of control animals, while the HS-deficient matrix was markedly softer.

The enhanced cartilage stiffness found in *Col2-rtTA-Cre;Ext1^e2fl/e2fl^*animals might indicate an increased stability of the cartilage against abrasion, in line with the reduced progression of OA in *Col2-rtTA-Cre;Ext1^e2fl/e2fl^ Ndst1^+/-^* and *Col2-Cre;Ndst1^fl/fl^*mutants (Severmann, 2020). Contradictingly, elevated AC stiffness has been associated with an accelerated onset of OA in mice lacking the ECM components Matn-4 (Li, 2020), the PG Decorin (Gronau, 2017) or carrying a hypomorphic allele of Can (Alberton, 2019). In contrast, our analysis showed elevated levels of Matn-3,-4 and Acan in the HS-deficient matrix, but a softened matrix has been linked to accelerated OA as well, as mice with a loss of Col6a1 exhibit a reduced YM of the pericellular matrix directly surrounding the chondrocytes and are prone to develop OA upon ageing (Alexopoulos, 2009). In line with pour findings in mutant mice with altered HS levels or modification patterns, *Sdc4^-/-^* animals are protected from OA progression as well (Echtermeyer, 2009). Of note, an increased level of CS has been detected in the nucleus pulposus of intervertebral discs of these mice (Sao, 2024). These findings highlight that HS play a key role for cartilage integrity and that ECM composition and stiffness need to be tightly regulated to ensure cartilage homeostasis.

We found increased levels of CS and its carrier proteins Acan and Pcan in the HS-deficient matrix of *Col2-rtTA-Cre;Ext1^e2fl/e2fl^* mice. This is in line with the excessive, non-stochiometric upregulation of CS in *Ext1^gt/gt^* mutants that we reported previously (Bachvarova, 2020). There are a few publications pointing to a role of CS in signalling regulation beyond its structural relevance (Willis, 2012) (Dyck, 2015). While the combined stiffening of the wildtype-like matrix and the softening of the HS-deficient matrix may not be sufficient to explain the protection from OA development observed in *Col2-rtTA-Cre;Ext1^e2fl/e2fl^* mice, a relevant factor likely is an altered bioavailability of essential signalling molecules for cartilage homeostasis, including members of the FGF, TGF, HH and WNT protein families that have been shown to interact with HS (Bishop, 2007) and CS (Schwartz, 2022). The CS-enriched matrix might sequester signalling molecules with catabolic function, such as Wnts. Unravelling whether a reduced bioavailability of catabolic factors or an enhanced release of anabolic factors from the HS-deficient matrix is the underlying mechanism of the protection observed in *Col2-rtTA-Cre;Ext1^e2fl/e2fl^* animals will require further investigation.

Another phenotype that has been linked to the loss of HS and has been studied in in *Col2-rtTA-Cre;Ext1^e2fl/e2fl^* mice is the genetic disorder Multiple Osteochondroma (MO; OMIM #133700) that develop upon clonal loss of HS in chondrocytes at the border of the GP. Increased BMP signalling has been hypothesised as one of the main drivers of MO (Huegel, 2013a) (Catheline, 2025) and we have previously demonstrated that HS-deficient chondroprogenitors from *Ext1^gt/gt^* embryos are more susceptible to Bmp stimulation (Gerstner, 2021). However, activation of Bmp signalling at the molecular level is not well understood. Interestingly, it has been shown that Integrin clustering in FA regulates TGFß and Bmp signalling by recruiting the respective receptors (Rys, 2015) (Sales, 2022). The clear upregulation of Integrin pathway components in HS-deficient clones of *Col2-rtTA-Cre;Ext1^e2fl/e2fl^* mice might point to a reorganisation of Integrin-containing cell-matrix contacts and thus enhanced clustering of Bmp-receptors. Alternatively, the bioavailability of Bmp signalling molecules might be enhanced in the HS-deficient context as outlined above.

Interestingly, while we detected increased levels of Integrin pathway in the HS-deficient clones, treatment of MEFs with the HS antagonist Surfen during cell adhesion showed impaired Integrin-dependent processes. This is in line with heparinase II treatment of primary bovine aortic endothelial cells leading to reduced cell adhesion after an incubation time of 30min (Moon, 2005) and the adhesion defects reported for ItgB1 deficient chondrocytes from *Col2-Cre;β1^fl/f^*^l^ mice (Aszodi, 2003), again highlighting a close relation between HS and Integrin function. Life cell imaging of the critical adhesion period from 0.5 to 1.5h post-seeding showed that more MEFs formed filopodia-like membrane protrusion upon Surfen treatment, similar to *in vivo* observations from *ext2-* and *ext3*-defcient zebrafish mutants where cells of the developing lateral line display an extensive formation of filopodia (Venero Galanternik, 2015). Under physiological conditions, initial adhesion complexes are formed in filopodia and filopodia with stable FA are converted to lamellipodia later on (Mattila, 2008). This process seemed to be disturbed in the Surfen-treated MEFs, which is substantiated by a reduced number of Surfen-treated cells that polarized and migrated during the timelapse recording. The described effects were largely compensated after 24h, highlighting a critical role of HS especially for early cells adhesion. HS on the cell surface might play a role in initiating cell-matrix contacts by electrostatic interaction between the negatively charged HS with positively charged surface of cell culture dishes *in vitro* or positively charged protein moieties of ECM components *in vivo* (Fig. 08). This notion is corroborated by the presence of HS-binding motifs in cartilage ECM components such as Collagens, Fibronectin and Laminins (Delacoux, 1998) (Nielsen, 2001) (Mahalingam, 2007). Still, treatment of already adherent MEFs led to an opposing effect and the addition Surfen resulted in an increased formation of FA and SF. This resembles the increased levels of Integrin pathway components detected in HS-deficient clusters of *Col2-rtTA-Cre;Ext1^e2fl/e2fl^*mice, where chondrocytes are permanently exposed to a loss of HS, and underlines the importance of considering the tissue context when studying Integrin function. Due to a lack of initial cell-matrix interactions upon loss of HS, the formation of stable cell-matrix adhesions might be delayed in the mutant chondrocytes. This signal could be interpreted as the matrix being too soft and induce a compensatory upregulation of matrix and Integrin pathway components.

**Figure 8:**
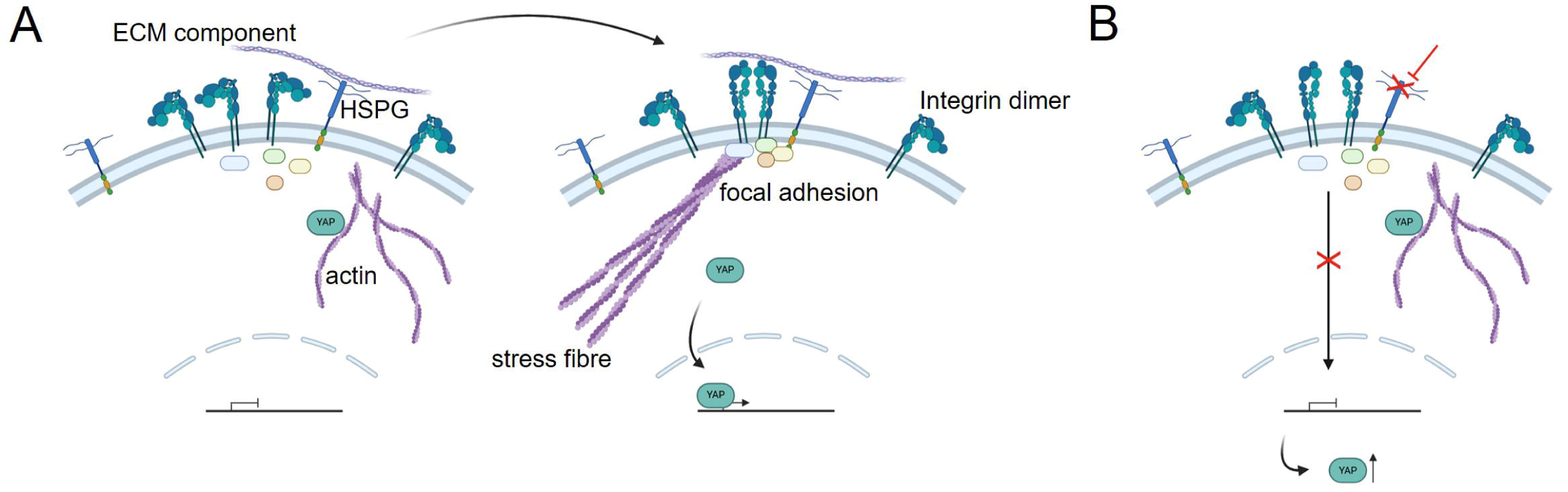
Cell surface HS are essential for cell-matrix interactions. Model of proposed working mechanism (Buchholz, 2026b). (A) Under physiological conditions, cell surface HS provide initial contacts with ECM components via HS-binding motifs. This promotes clustering of Integrin receptors contributing to the formation of stable FA complexes. Signalling via Integrin-dependent cascades, including YAP, maintain the status of the cell. (B) Upon loss of HS function, initial cell-matrix interactions are impaired, interfering with the formation of FA and SF. To rescue the inhibited cell-matrix adhesion, production of Integrin pathway components and downstream factors, such as YAP, are upregulated.

### HS control GAG synthesis via subcellular localisation of YAP

A major signalling cascade transducing information on matrix stiffness into the nucleus is the hippo pathway via YAP. The localisation of YAP is controlled by intracellular tension generated by the actin cytoskeleton, including binding of YAP to filamentous actin in the cell periphery and the stretching of nuclear pores (Seo, 2018). In particular, the formation of SF is orchestrated with the subcellular localisation of YAP (Dupont, 2011) (Heng, 2021). We found that in MEFs treated with Surfen during adhesion less SF are formed, and YAP is primarily found in the cytoplasm. Similarly, YAP shifts to the cytoplasm when primary rat chondrocytes are seeded onto a soft rather than a hard surface (Zhong, 2013), indicating that impaired HS function might lead cells to wrongly perceive the substrate as soft due to impaired cell-matrix adhesion. While the disruption of FA and SF formation by Surfen was recovered after 24h, YAP remained in the cytoplasm at this time point, and we did not detect a shift of YAP in MEFs treated with Surfen for 1h after adhesion, pointing to a slower response of YAP localisation to HS-dependent cues compared to FA and SF formation.

Still, the level of YAP was clearly upregulated in *in vivo*, where chondrocytes are permanently HS-deficient. As outlined above, HSPGs control the bioavailability of signalling molecules, the organisation of the cytoskeleton, and the signalling transduction downstream of Integrins. All these processes are also involved in the regulation of YAP localisation and activity. *Vice versa*, evidence for the regulation of HSPGs levels via YAP is scarce: A genetically modified melanoma cell line with constitutive active YAP showed enhanced cell surface levels of HS (Dieter, 2022) and YAP seemed to be involved in the regulation of Prg4 expression downstream of Cdc42 (Delve, 2020).

We showed that YAP activation with LPA during chondrogenic differentiation of micromass cultures increased the deposition of GAGs, in line with the increased levels of CS and PG core proteins detected in HS-deficient clusters of *Col2-rtTA-Cre;Ext1^e2fl/e2fl^* mice. This shows that YAP acts upstream of GAG synthesis and points to a positive effect of YAP activation on ECM synthesis. It has previously been reported that YAP1 levels are reduced in endplate cartilage from patients with degenerated intervertebral discs, and that the loss of COL2A1 and ACAN observed in those cells was rescued by viral overexpression of YAP (Ding, 2022). Likewise, YAP inhibition in primary human chondrocytes using VP reduced the expression of the anabolic genes SOX9, COL2A1 and ACAN, while YAP activation using LPA enhanced COL2A1 expression (Cui, 2023), again supporting an anabolic role of YAP for cartilage ECM. Contradictingly, it has been reported that simultaneous knock down of Yap (Ding, 2022) and Taz using siRNA in ATDC5 cells increases the mRNA expression of chondrogenic markers, such as Sox9, Col2a1 and Acan, and that LPA decreased their expression (Hallstrom, 2023), indicating a negative effect of YAP on ECM synthesis. This observation is limited, though, by the use of a murine teratoma cell line rather than primary human chondrocytes.

We showed that Surfen-treatment enhanced GAG production in line with existing literature (Huegel, 2013b) and that activation of YAP by LPA resulted in the same effect. YAP inhibition using VP mitigated the effect of Surfen, but did not entirely abolish it, suggesting that HS-function-dependent GAGs synthesis is transduced not only through YAP but also through other parallel mechanisms.

Taken together, we conclude that cell surface HSPGs are essential for the control of ECM composition and thus the mechanical properties of cartilage matrix. We confirm that HS are vital for Integrin-dependent processes, such as cell adhesion, polarisation and migration, and show that the localisation of YAP is not only regulated by HS, but that YAP also controls the synthesis of GAGs. When HS are lost in the context of cartilage tissue, this likely leads to the formation of less functional cell-matrix interactions and an attempt to compensate the lack of adhesion by upregulating the synthesis of specific matrix and Integrin pathway components. A change in matrix composition, stiffness and biological function might also affect the tissue homeostasis in degenerative joint disease, such as OA, and the formation of osteochondromas in MO.

