## Supplementary Table 01 for "Heparan sulfate-function is essential for Integrin-dependent cell-matrix interactions and regulates glycosaminoglycan synthesis synthesis through YAP"

| **Allele** | **Primer 1** | **Primer 2** | **Primer 3** | **Annealing**  **[°C]** |
| --- | --- | --- | --- | --- |
| *Col2-rtTA-Cre* | gagtgatgaggttcgcaaga | ctacaccagagacgg |  | 55 |
| *Ext1^e2fl/e2fl^* | gagtccatcctgctctgcat | ttgttgcatgggaaagacaa |  | 62 |
| *Ext1^fl/fl^* | ggagtgtggatgagttgaag | caacactttcagctccagtc | cctgagaagcccaagctca | 62 |
| *R26R-LacZ* | aagaccgcgaagagtttgtc | aaagtcgctctgagttgttat | ggagcgggagaaatggatatg | 58 |
| *R26R-mTmG* | ctctgctgcctcctggcttct | cgaggcggatcacaagcaata | tcaatgggcgggggtcgtt | 60 |

**Supplementary Table 1: Primers for genotyping PCRs**
