## Supplementary File 01 for "Heparan sulfate-function is essential for Integrin-dependent cell-matrix interactions and regulates glycosaminoglycan synthesis synthesis through YAP"

### 1 Statistical models

For all models, the index $i$ denotes individual observations but is omitted where possible for improved readability.

#### 1.1 Fig. 4B: Numbers of MEFs adhering to the cell culture dish upon Surfen treatment

$$
\begin{aligned}
y_i &\sim \text{NegBin}\left(\mu_i,\ \phi_{c_i,t_i}\right) \\
\log\left(\mu_i\right) &= \alpha_{treat_i,time_i} + \gamma_{e_i} \\
\alpha_{treat,time} &\sim \text{Normal}\left(3,\ 3\right) \\
\gamma_e &\sim \text{Normal}\left(0,\ 1\right), \quad \sum_{e=1}^{2} \gamma_e = 0 \\
\phi_{c,t} &\sim \text{LogNormal}\left(0,\ \sigma_\phi\right) \\
\sigma_\phi &\sim \text{Gamma}\left(2,\ 2\right)
\end{aligned} \tag{1}
$$

We modelled the number of cells $y_i$ with a negative binomial distribution parameterized by a mean and an overdispersion term. Both distributional parameters depend on the interaction between the treatment levels $treat \in 1, ..., 5$ and the time points $time \in 1, ..., 5$. The experiment effect $\gamma_e$, $e \in 1, 2$ is treated as a deviation from the overall mean with a sum-to-zero constraint. We expressed effects as ratios of the expectation values for specific treatment levels and time points, e.g. $\delta = \frac{\mu_{treat_1,time_1}}{\mu_{treat_2,time_2}} = \exp\left(\alpha_{treat_1,time_1} - \alpha_{treat_2,time_2}\right)$, with the corresponding $\pi$-value computed as $\pi(\delta, 1)$.

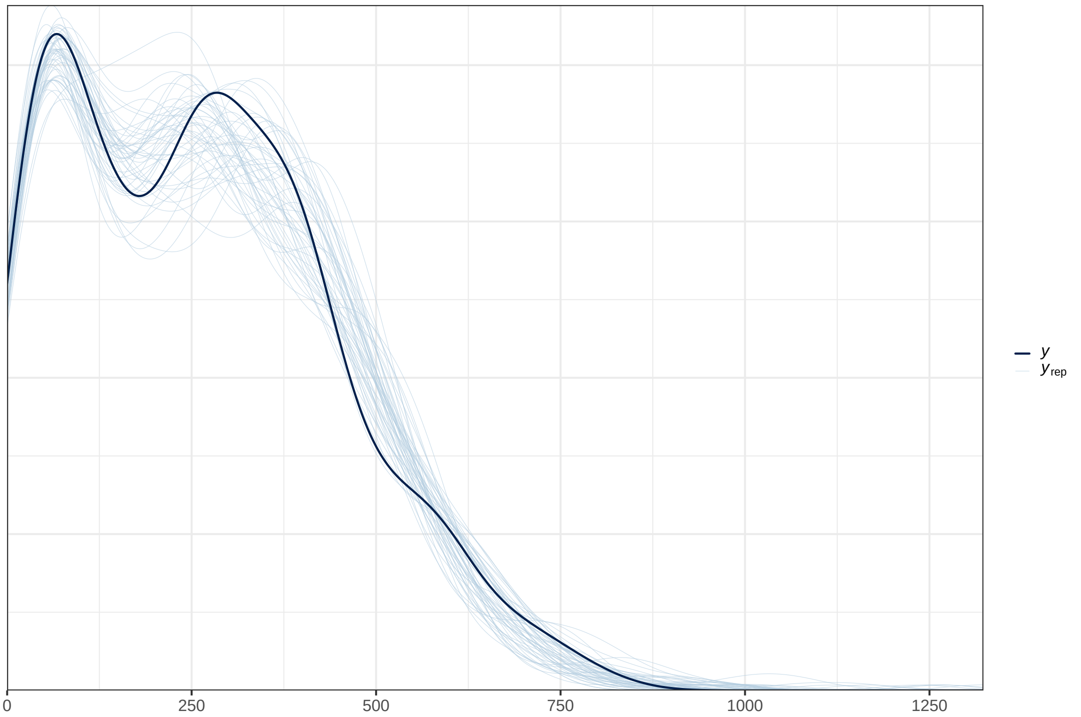


Figure 1: Posterior predictive check for the numbers of adhering MEFs. Density of the observed data is shown as the thick line, replicated data generated from the posterior are shown in thin lines.

#### 1.2 Fig. 4G-J: Life cell imaging

##### 1.2.1 Fig. 4G, H: Membrane phenotypes of MEFs at 0.5 and at 1.5h post seeding upon Surfen treatment

$$\begin{aligned} y_i &\sim \text{BetaBin}(\text{N}_i,\, p_i \cdot \phi,\, (1-p_i)\cdot \phi) \\ \text{logit}(p_i) &= \alpha_{\text{type}_i,\,\text{treat}_i,\,\text{time}_i} \\ \alpha_{\text{type, treat, time}} &\sim \text{Normal}(0,\, 2) \\ \phi &\sim \text{Logormal}(0,\, 1) \end{aligned} \tag{2}$$

To account for overdispersion, we modelled the number of cells $y_i$ that forms either pseudopodia, lamellipodia, or filopodia (type) using a beta-binomial distribution. The probability $p_i$ is parameterized on the logit scale and depends on the interaction between type, treatment (control, Surfen) and time point (start, end). Effects are expressed as differences in probabilities between combinations of type, treatment, and time points, e.g. $\delta = p_{type_1,treat_1,time_1} - p_{type_2,treat_2,time_2}$, with the corresponding $\pi$-value computed as $\pi(\delta, 0)$.

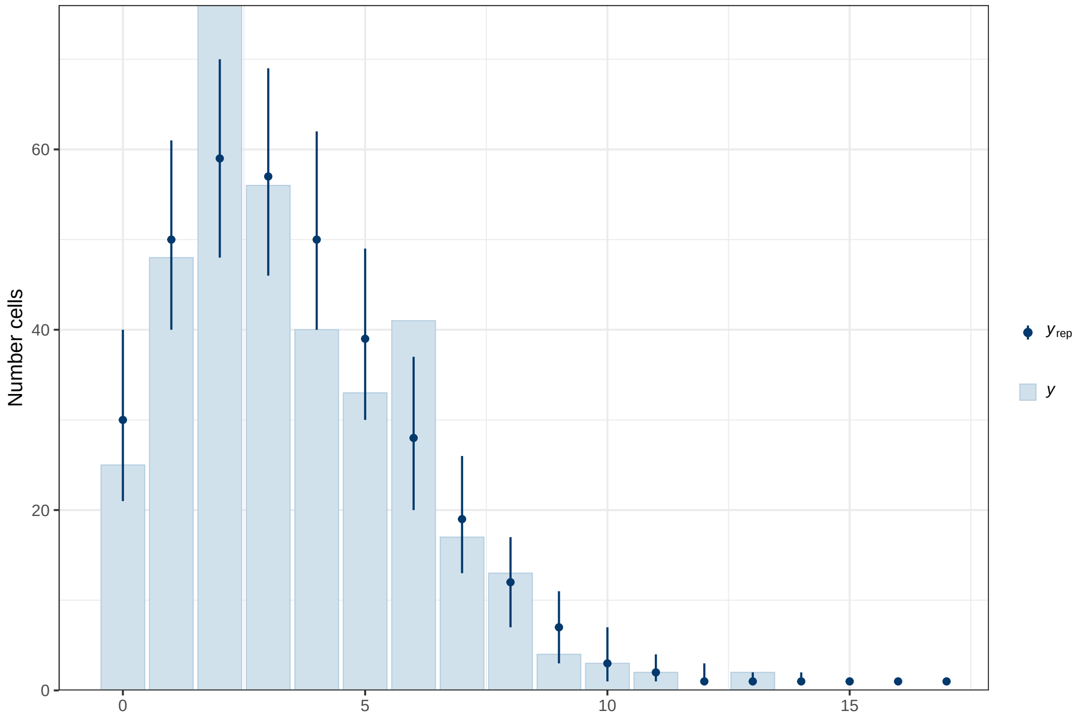


Figure 2: Posterior predictive check for the analysis of membrane phenotypes at 0.5 and at 1.5h post seeding. The histogram shows the observed number of cells forming a defined mebrane phenotype. Model predictions are summarized by the posterior median (dot) and the 90% credible interval (bar).

###### 1.2.2 Fig. 4I: Numbers of filopodia per cell at 0.5 and at 1.5h post seeding upon Surfen treatment

$$\begin{aligned} y_i &\sim \text{NegBin}\left(\mu_i,\, \phi\right)^T [1, \infty) \\ \log\left(\mu_i\right) &= \alpha_{time_i, treat_i} + \gamma_{e_i} \\ \alpha_{time,treat} &\sim \text{Normal}\left(0,\, 2\right) \\ \gamma_e &\sim \text{Normal}\left(0,\, 1\right), \quad \sum_{e=1}^{3} \gamma_e = 0 \\ \phi &\sim \text{LogNormal}\left(0,\, 2\right) \end{aligned} \tag{3}$$

We modelled the number of filopodia per cell $y_i$ using a negative binomial distribution truncated at 1 – since we are only interested in cells that form at least one filopodium – and parameterized by a mean and an overdispersion term. The expectation value $\mu$ depends on the interaction between the treatment (control, Surfen) and time point (t = 0.5h, t = 1.5h). The experiment effect $\gamma_e$, $e \in 1, 2, 3$ is treated as a deviation from the overall mean with a sum-to-zero constraint. We expressed effects as ratios of the expected values for combinations of treatment and time points, e.g. $\delta = \exp\left(\alpha_{time_1,treat_1} - \alpha_{time_2,treat_2}\right)$, with the corresponding $\pi$-value computed as $\pi(\delta, 1)$.

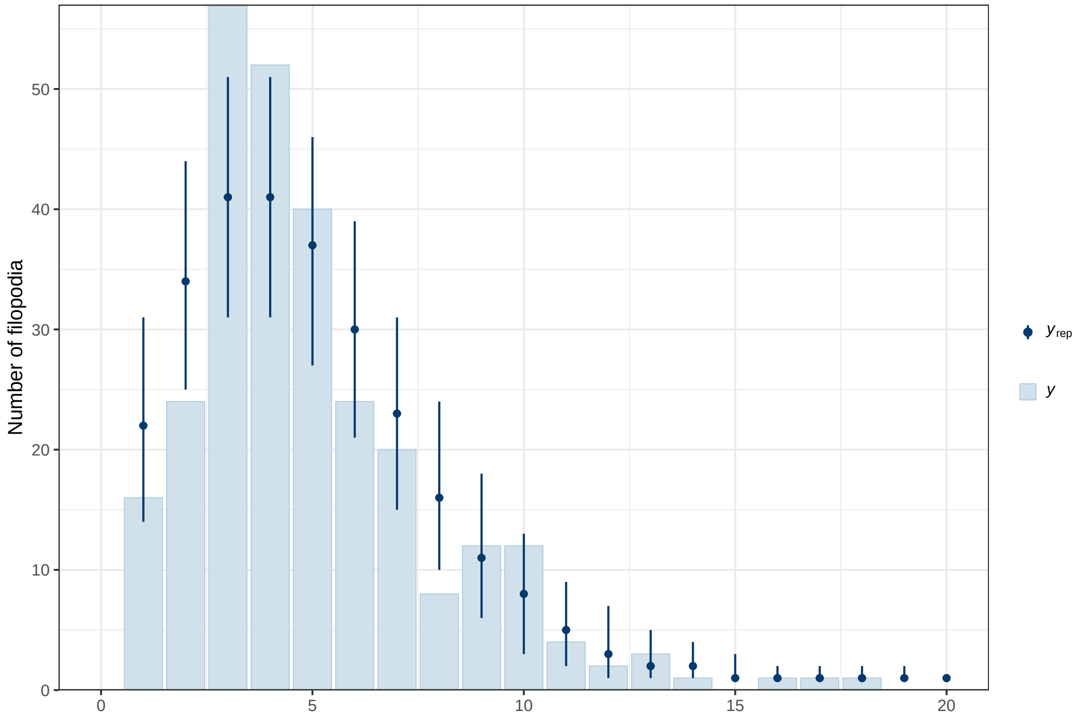


Figure 3: Posterior predictive check for the analysis of filopodia-like membrane protrusions at 0.5 and at 1.5h post seeding. The histogram shows the observed number of filopodia per cell. Model predictions are summarized by the posterior median (dot) and the 90% credible interval (bar).

##### 1.2.3 Fig. 4J: Polarisation and migration of MEFs from 0.5 to 1.5h post seeding upon Surfen treatment

$$\begin{aligned} y_i &\sim \text{Binomial}(\text{N}_i,\, p_i) \\ \text{logit}(p_i) &= \alpha_{\text{type}_i,\,\text{treat}_i} \\ \alpha_{\text{type, treat}} &\sim \text{Normal}(0,\, 2) \end{aligned} \tag{4}$$

The number of polarising or migrating cells $y_i$ is modelled using a binomial distribution with success probability $p_i$ and total number of cells $N$. The probability $p_i$ is parameterized on the logit scale and depends on the interaction between cell type (polarising or migrating) and treatment (control or Surfen). The experiment effect $\gamma_e$, $e \in 1, 3$ is treated as a deviation from the overall mean with a sum-to-zero constraint. Effects are expressed as differences of probability for treatment and time points, e.g. $\delta = p_{type_1,treat_1} - p_{type_2,treat_2}$, with the corresponding $\pi$-value computed as $\pi(\delta, 0)$.

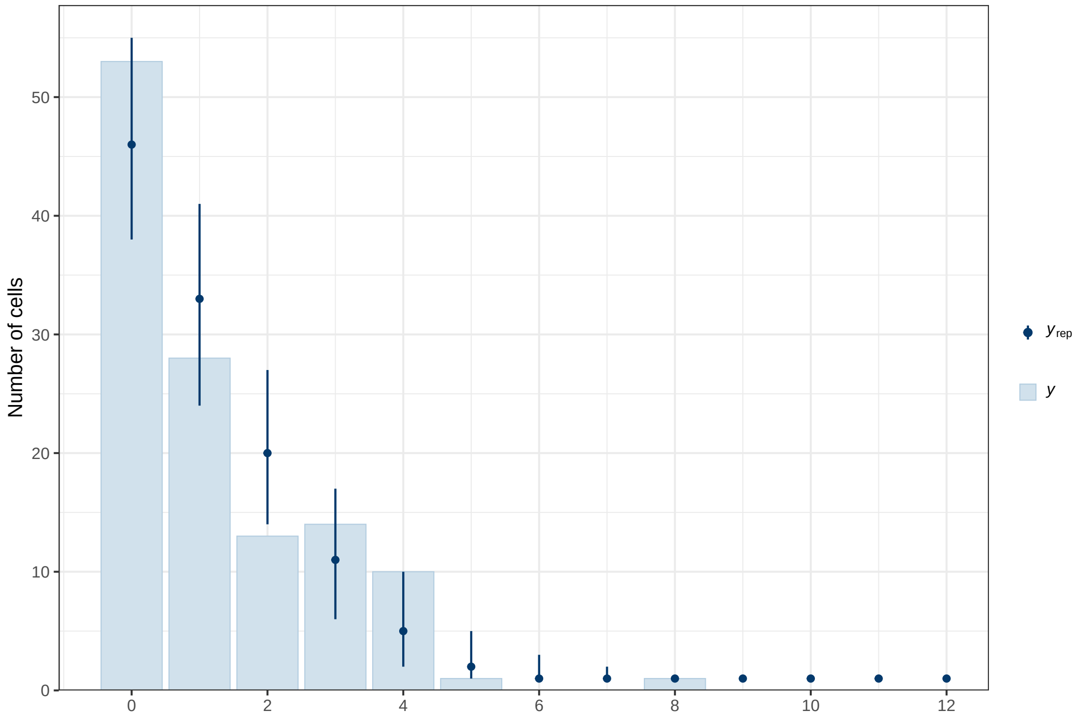


Figure 4: Posterior predictive check for the analysis of time lapse videos from 0.5 to 1.5h post seeding. The histogram shows the observed numbers of migrating and polarising cells. Model predictions are summarized by the posterior median (dot) and the 90% credible interval (bar).

##### 1.3 Fig. 5H: Ellipticity of CHO wildtype and pgsD-677 cells at 1.25h post seeding

$$
\begin{aligned}
y_i &\sim \theta_{type_i,surface_i} \cdot \mathrm{BetaProp}(p_i^{low},\, \phi^{low}_{type_i,surface_i}) + \\
&\quad (1-\theta_{type_i,surface_i}) \cdot \mathrm{BetaProp}(p_i^{high},\, \phi^{high}_{type_i,surface_i}) \\
\log(\delta) &= \alpha^{\delta} + \beta^{\delta}_{type} + \beta^{\delta}_{surface} + \beta^{\delta}_{rep} \\
\mathrm{logit}(p^{high}) &= \alpha + \beta_{type} + \beta_{surface} + \beta_{rep} + \delta \\
\theta &\sim \mathrm{Beta}(1,\, 1) \\
\alpha, \beta_{type}, \beta_{surface}, \beta_{rep} &\sim \mathrm{Normal}(0,\, 0.3) \\
\alpha^{\delta}, \beta^{\delta}_{type}, \beta^{\delta}_{surface}, \beta^{\delta}_{rep} &\sim \mathrm{Normal}(0,\, 0.1) \\
\phi^{low}, \phi^{high} &\sim \mathrm{Normal}(\mu_{\phi},\, \sigma_{\phi}) \\
\mu_{\phi} &\sim \mathrm{Normal}(0,\, 50) \\
\sigma_{\phi} &\sim \mathrm{Normal}(1,\, 1)
\end{aligned}
\tag{5}
$$

The ellipticity $y_i$ of cells $\left(\frac{shortaxis}{longaxis}\right)$ is modelled using a mixture of two beta distributions, motivated by the bimodality observed in the data (round cells with $p^{high}$ and elongated cells with $p^{low}$) across different cell types and surfaces. We used the beta-proportion parameterization, where the beta distribution is expressed in terms of an expectation parameter $p$ and an overdispersion parameter $\phi$. The mixing weight $\theta$, which determines the contribution of each component, depends on the interaction between cell type (pgsD-677, wild type) and surface (RGD, RGD+iHS). The expectation values of both components, $p^{high}$ and $p^{low}$, are parameterized on the logit scale and depend on cell type, surface, and biological replicate. All effects are coded as centered deviations (sum-to-zero constraints), so that $\alpha$ represents the overall intercept. Effects are expressed as differences in the overall expected probability – i.e., the mixture expectation across both components – for combinations of cell type and surface, e.g. $\delta = \theta_{type_1,surface_1} \cdot p^{low}_{type_1,surface_1} + (1 - \theta_{type_1,surface_1}) \cdot p^{low}_{type_1,surface_1} - \theta_{type_2,surface_2} \cdot p^{low}_{type_2,surface_2} + (1 - \theta_{type_2,surface_2}) \cdot p^{low}_{type_2,surface_2}$, with the corresponding $\pi$-value computed as $\pi(\delta, 0)$.

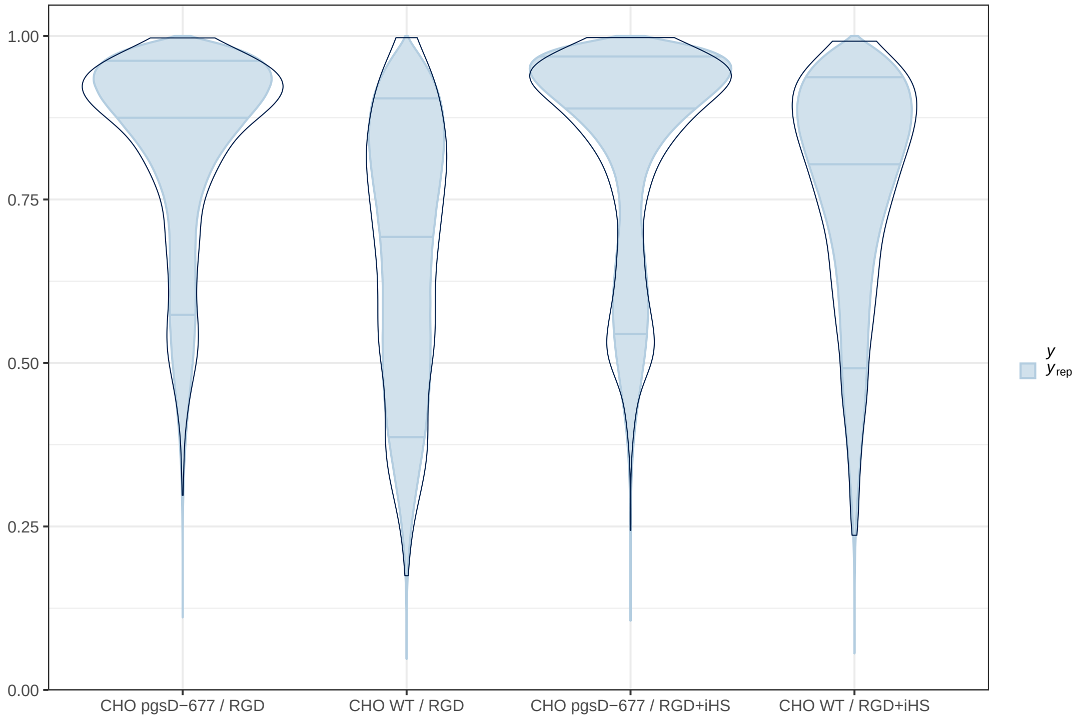


Figure 5: Posterior predictive check for analysis of the ellipticity of CHO WT and CHO pgsD-677 cells at 1.25h post seeding. The plot shows the ellipticity defined as the ration of short cell axis / long cell axis. Density of the observed data is shown as violin plots (thick line) for each combination of cell type and surface. Densities of replicated data generated from the posterior are shown as filled violins.

##### 1.4 Fig. 6B, D, F, G: Numbers of MEFs with specific IF staining patterns upon Surfen treatment

$$
\begin{aligned}
y_i &\sim \text{Binomial}(\text{N}_i,\, p_i) \\
\text{logit}(p_i) &= \alpha_{\text{cond}_i,\,\text{treat}_i} + \gamma_e \\
\alpha_{\text{cond},\,\text{treat}} &\sim \text{Normal}(0,\, 2) \\
\gamma_e &\sim \text{Normal}(0,\, 1), \quad \sum_{e=1}^{3} \gamma_e = 0
\end{aligned}
\tag{6}
$$

We applied the same statistical model to all three fluorescence staining data sets, where $y_i$ denotes the number of cells showing the respective indicator: focal adhesions in the immunofluorescence against Paxillin data set, stress fibres in the fluorescence-labelled Phalloidin data set, and nuclear signals in the immunosfluorescence against YAP data set. The number of cells $y_i$ is modelled using a binomial distribution with success probability $p_i$ and total number of cells $N$. The probability $p_i$ is parameterized on the logit scale and depends on the interaction between condition ('1h treatment during cell adhesion', '24h treatment during cell adhesion', ' 1h treatment after cell adhesion') and treatment (control, Surfen). The experiment effect $\gamma_e$, $e \in 1, 2, 3$ is treated as a deviation from the overall mean with a sum-to-zero constraint. Effects are expressed as differences of probability for treatment and condition, e.g. $\delta = p_{cond_1,treat_1} - p_{cond_2,treat_2}$, with the corresponding $\pi$-value computed as $\pi(\delta, 0)$.

###### 1.4.1 Fig. 6B, G: Proportion of cells forming focal adhesions (FA)

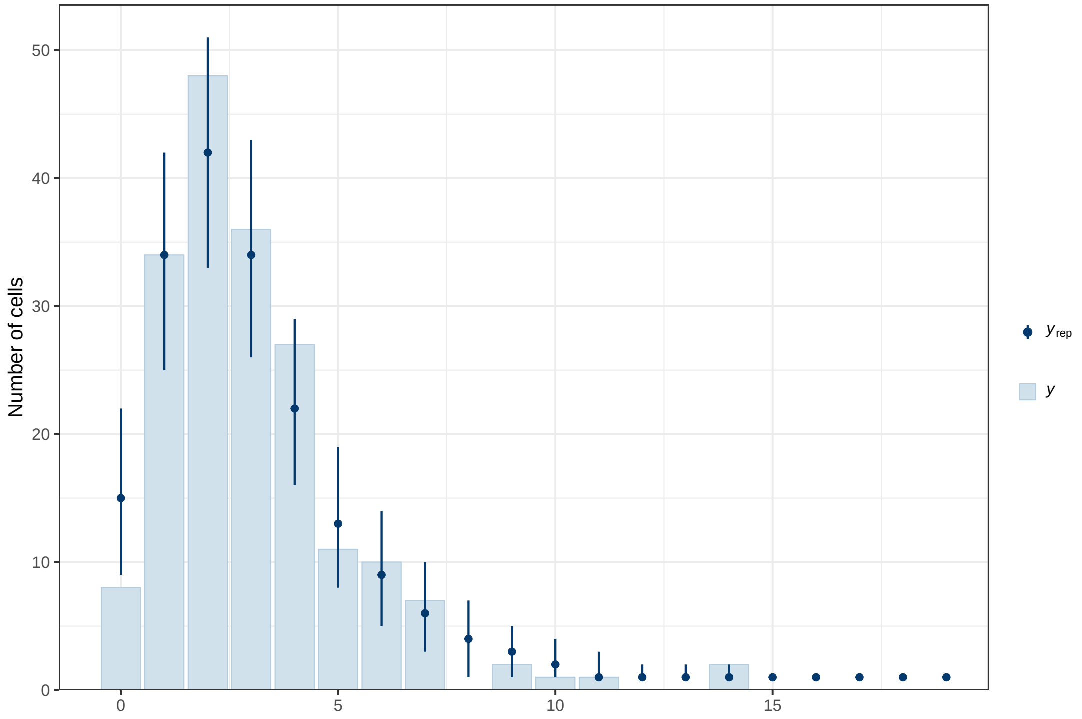


Figure 6: Posterior predictive check for the detection of FA by immunofluorescence staining of Paxillin. The histogram shows the observed number of cells forming focal adhesions. Model predictions are summarized by the posterior median (dot) and the 90% credible interval (bar).

##### 1.4.2 Fig. 6D, G: Proportion of cells forming stress fibres (SF)

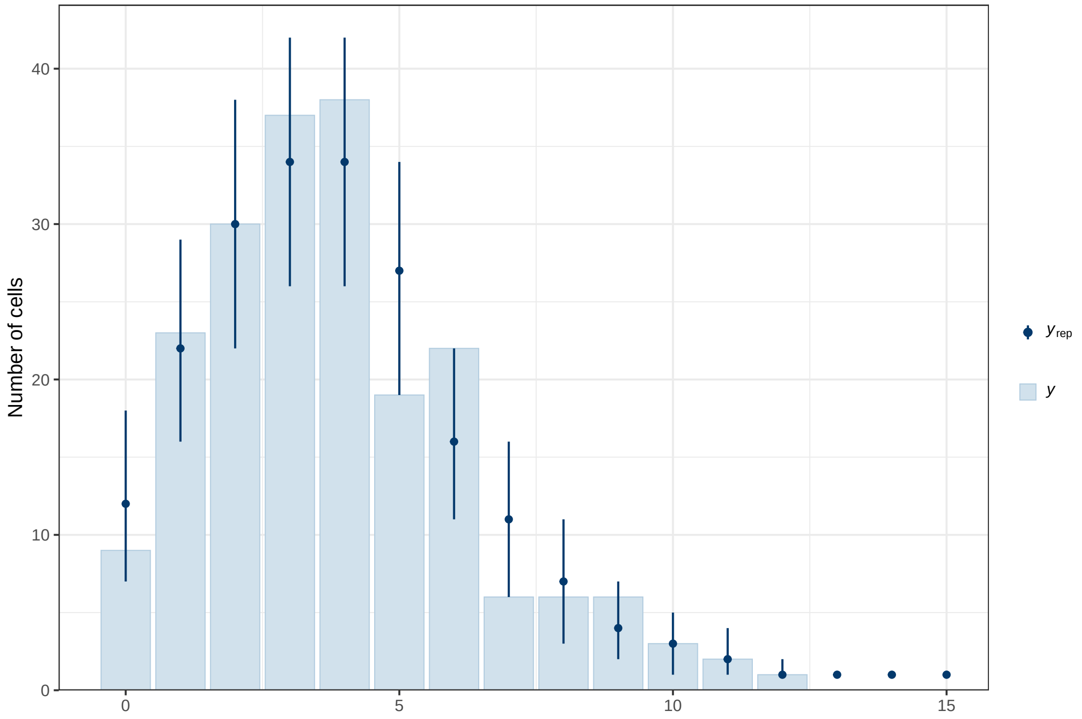


Figure 7: Posterior predictive check for the detection of SF by staining of the actin cytoskeleton using a fluorescence-labelled Phalloidin. The histogram shows the observed number of cells forming stress fibres. Model predictions are summarized by the posterior median (dot) and the 90% credible interval (bar).

##### 1.4.3 Fig. 6F, G: Proportion of cells showing a clear nuclear YAP (nYAP) signal

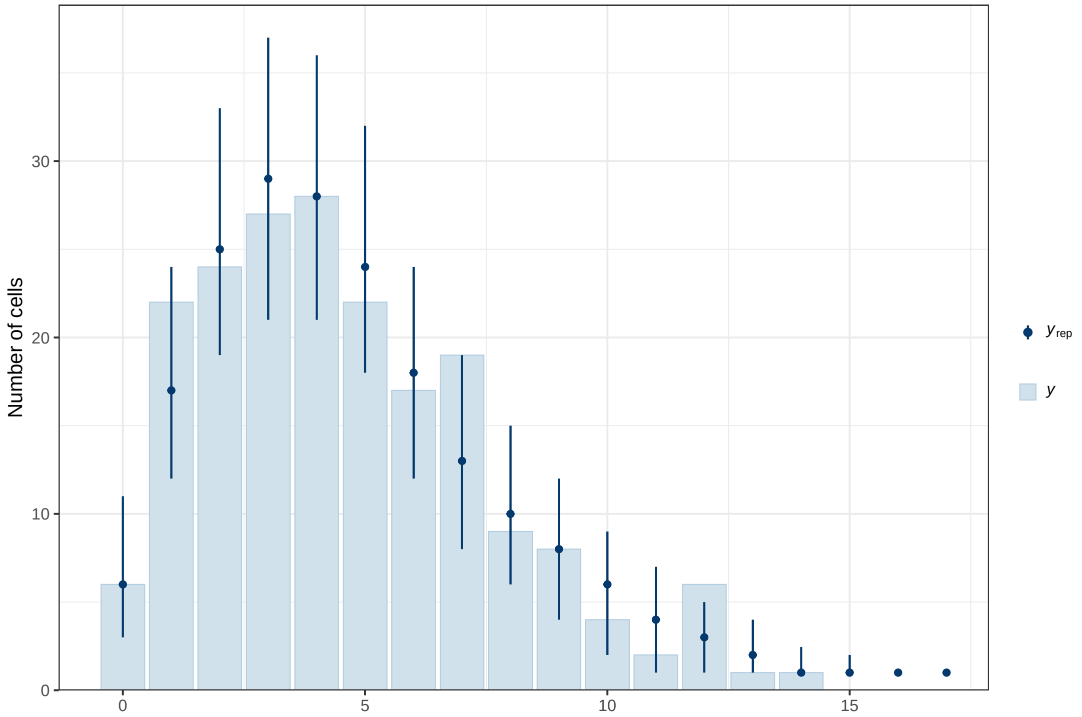


Figure 8: Posterior predictive check for the detection of nYAP by immunofluorescence. The histogram shows the observed number of cells showing nuclear signals. Model predictions are summarized by the posterior median (dot) and the 90% credible interval (bar).

#### 1.5 Fig. 7: Relative absorbance of solubilised Alcian blue from micromass culture upon pharmacological treatments

$$\begin{aligned} y_i &\sim \text{BetaProp}(p_{treat_i,rep_i},\, \phi) \\ p_{treat,rep} &\sim \text{BetaProp}(p_{treat},\, \phi') \\ p_{treat} &\sim \text{Beta}(1,\, 1) \\ \phi,\, \phi' &\sim \text{Lognormal}\,(0,\, 2) \end{aligned} \tag{7}$$

We applied the same statistical model to all three data sets, where $y_i$ denotes the relative absorbance of solubilised Alcian blue, defined as $\frac{I}{I_0}$, with $I$ being the transmitted light intensity and $I_0 > I$ the incident light intensity. The relative absorbance $y_i$ is modelled using a beta-proportion distribution with the expected probability $p_{treat,rep}$ (depending on treatment and replicate) and an overdispersion parameter $\phi$. To account for biological variability among replicates, we used a hierarchical model in which each treatment has an overall expected probability $p_{treat}$, from which the replicate-level parameters $p_{treat,rep}$ are drawn. Effects are expressed as differences of probability for treatments, e.g. $\delta = p_{treat_1} - p_{treat_2}$, with the corresponding $\pi$-value computed as $\pi(\delta, 0)$.

##### 1.5.1 Fig. 7C: Relative GAG content upon YAP activation

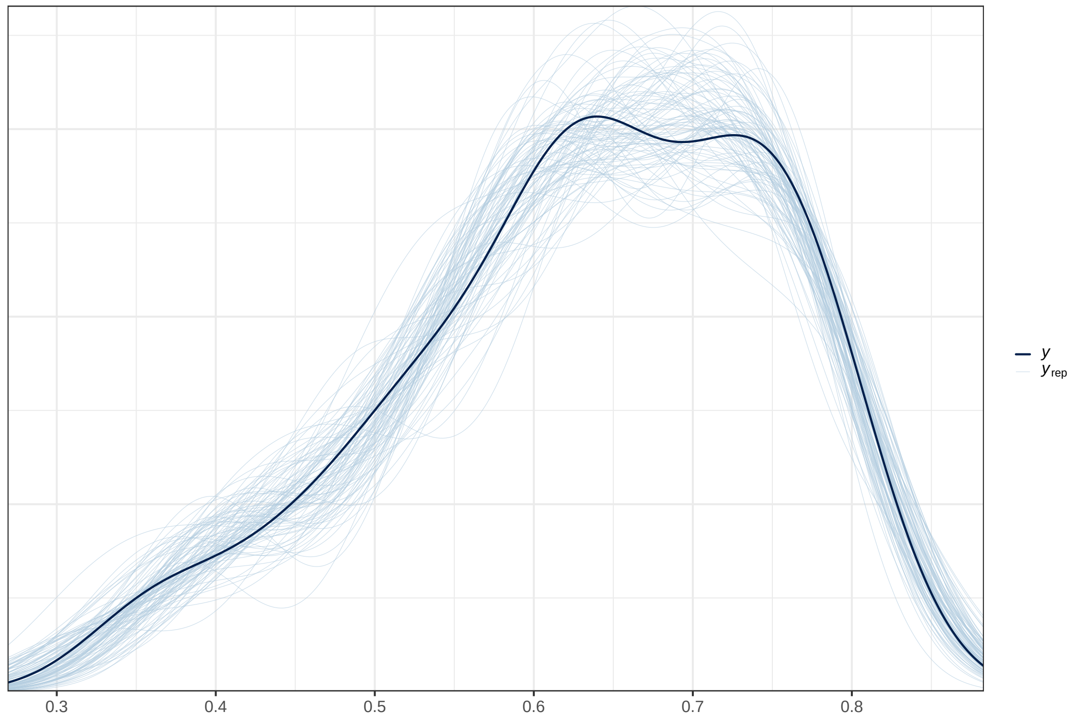


Figure 9: Posterior predictive check for the treatment of micromass cultures with the YAP activator lysophosphatidic acid (LPA). Density of the observed data is shown as the thick line, replicated data generated from the posterior are shown in thin lines.

##### 1.5.2 Fig. 7E: Relative GAG content upon YAP inhibition

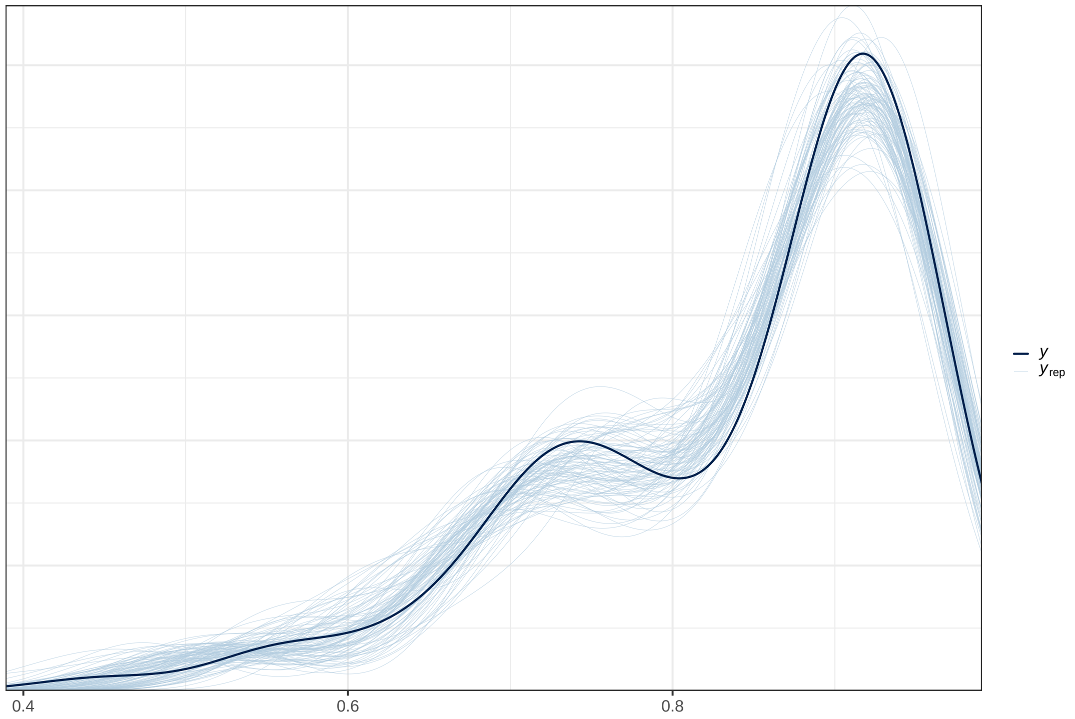


Figure 10: Posterior predictive check for the treatment of micromass cultures with the YAP inhibitior Verteporfin (VP). Density of the observed data is shown as the thick line, replicated data generated from the posterior are shown in thin lines.

##### 1.5.3 Fig. 7G: Relative GAG content upon combined HS and YAP inhibition

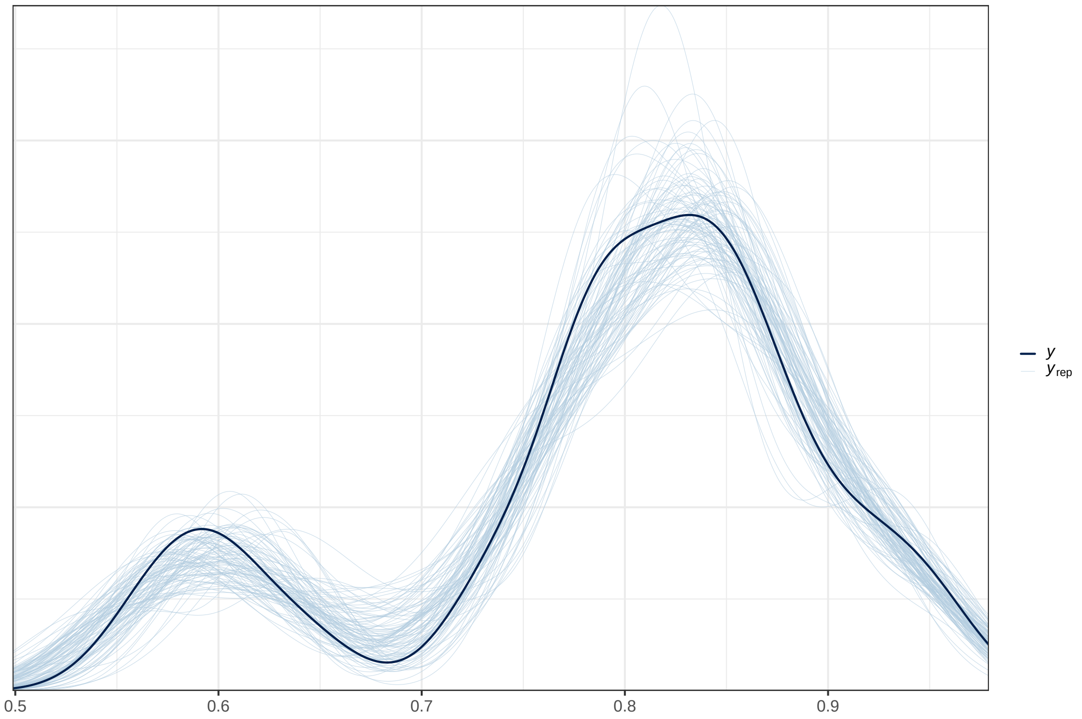


Figure 11: Posterior predictive check for the combined treatment of micromass cultures with the HS antagonist Surfen and the YAP inhibitior VP. Density of the observed data is shown as the thick line, replicated data generated from the posterior are shown in thin lines.
